# Directed insulin secretion from beta cells occurs at cortical sites devoid of microtubules at the edges of ELKS/LL5β patches

**DOI:** 10.1101/2024.10.31.621333

**Authors:** Margret Fye, Pranoy Sangowdar, Anissa Jayathilake, Pi’ilani Noguchi, Guoqiang Gu, Irina Kaverina

## Abstract

To maintain normal blood glucose levels, pancreatic beta cells secrete insulin into the bloodstream at specialized regions at the cell periphery, often called secretion hot spots. While many secretory machinery components are located all over the cell membrane, directed secretion relies on distinct cortical patches of the scaffolding protein ELKS and the microtubule (MT)-anchoring protein LL5β. However, using TIRF microscopy of intact mouse islets to precisely localize secretion events within ELKS/LL5β patches, we now show that secretion is restricted to only 5% of ELKS/LL5β patch area. Moreover, the majority of secretion occurs at the margins of ELKS patches. This suggests that additional factor(s) must be responsible for hot spot definition. Because the MT cytoskeleton plays a regulatory role in the insulin secretion process via both delivery and removal of secretory granules from the secretion sites, we test whether local MT organization defines secretory activity at hot spots. We find that the majority of secretion events occur at regions devoid of MTs. Based on our findings, we present a model in which local MT disassembly and optimal ELKS content are strong predictors of directed insulin secretion.

**Significance Statement:**

- Insulin has to be secreted directly into the bloodstream for efficient regulation of glucose metabolism. Molecular requirements for precise secretion location and microtubule-mediated regulation in directed secretion are uncharacterized.
- Using intact mouse islets and a live insulin secretion assay, we demonstrate that cortical patches containing ELKS and LL5β display secretory heterogeneity and secretion hot spots are precisely localized to patch edges that are devoid of microtubules.
- These findings suggest that microtubule absence is critical for secretion and secretion occurs away from presumed sites of secretory granule delivery. This expands the current knowledge of regulation and spatial characteristics of insulin secretion.

## Introduction

Pancreatic beta cells, which comprise the majority of exocrine pancreatic islets, secrete insulin in order to regulate blood glucose levels. When blood glucose levels rise, beta cells respond by secreting insulin to promoting glucose uptake in peripheral tissues. In the last decade, it has become clear that secretion occurs within specific subcellular regions in beta cells, often called secretion hot spots (Fu et al., 2019; Trogden et al., 2021). Moreover, several groups have characterized beta cells as having polarity with a defined basal side, which allows for directed secretion toward the vasculature (Bonner-Weir, 1988; Granot et al., 2009; Geron et al., 2015; Gan et al., 2017; Cottle et al., 2021; Jevon et al., 2024). In particular, secretion hot spots in beta cells are localized to areas of contact with vascular extracellular matrix, resulting in preferential insulin secretion into the bloodstream (Gan et al., 2018; Jevon et al., 2022). This directed secretion phenomenon is impaired under diabetic conditions (Ohara-Imaizumi et al., 2005; Almaça et al., 2015; Ohara-Imaizumi et al., 2019b; Jevon et al., 2024), making it a potential candidate for diabetes therapeutics.

Mechanistically, directed secretion is promoted by an elaborate molecular platform (Fye and Kaverina, 2023), initiated by transmembrane receptor integrins, especially integrin β1 after binding with their ligands within vascular extracellular matrix (ECM) (particularly, components of basement membrane produced by endothelial cells, laminin and collagen IV) (Kaido et al., 2004; Spelios et al., 2015; Gan et al., 2018; Jevon et al., 2022). This molecular platform includes a variety of components of focal adhesions (Rondas et al., 2011; Arous et al., 2013; van der Vaart et al., 2013; Bouchet et al., 2016), ion channels (Ohara-Imaizumi et al., 2019b; Jevon et al., 2022), exocytic machinery (Yasuda et al., 2010; Low et al., 2014), and proteins common to the active zone such as Munc18c and liprins (Oh and Thurmond, 2009; Low et al., 2014; Gan et al., 2017; Cottle et al., 2021). The molecular components of these platforms are characterized by precise spatial distribution, with the active zone proteins concentrated outside focal adhesion plaques (Lansbergen et al., 2006; Hotta et al., 2010; Bouchet et al., 2016; Fourriere et al., 2019; Noordstra et al., 2022). Interestingly, prior studies in other cell types locate exocytic events to the focal adhesion periphery (Grigoriev et al., 2007; Stehbens et al., 2014; Fourriere et al., 2019; Ren et al., 2025). However, detection of precise localization of insulin secretion events in relation to the active zone protein accumulation have been challenging for technical reasons and remain unclear to date.

One of the primary proteins responsible for directed insulin secretion is ELKS, so named for those residues enriched in its amino acid sequence. Several groups in the past decade or so have shown that ELKS plays a significant role in vascular-directed insulin secretion (Ohara-Imaizumi et al., 2005; Low et al., 2014; Ohara-Imaizumi et al., 2019b). ELKS is a scaffolding protein and functions to both bind several directed secretion proteins via its coiled-coil domains (Held et al., 2016, Held and Kaeser, 2018) as well as regulate voltage-dependent Ca^2+^ channel (VDCCs) to promote secretion (Ohara-Imaizumi et al., 2019a; Ohara-Imaizumi et al., 2019b). Knocking out elks is also known to impair directed insulin secretion, glucose tolerance, and Ca^2+^ flux (Ohara-Imaizumi et al., 2019b), making it one of the most critical components of directed insulin secretion.

ELKS’ binding partner LL5β is also known to associate with these directed secretion protein complexes (Lansbergen et al., 2006; Noordstra et al., 2022) and is known to bind microtubule (MT) plus ends via the MT end-binding protein CLASP2 (Lansbergen et al., 2006;, Basu et al., 2015). LL5β in other cell types is known to be an important part of the cortical MT stabilizing complex (CMSC) and is partly responsible for MT anchoring at the membrane (Hotta et al., 2010; van der Vaart et al., 2013; Noordstra et al., 2022) also recently showed that LL5β is required for clustering of insulin granule (IG) docking complexes.

The connection of directed secretion to MT function has been highlighted in our recent studies. Our group has extensively characterized the role of MTs in pancreatic beta cells. Historical thought was that the MT cytoskeleton in beta cells was arranged in an astral array, akin to fibroblasts, that allowed MT plus-end-directed trafficking of IGs to the periphery of the cell (Boyd et al., 1982, Varadi et al., 2003). While this plus-end-trafficking is important, our group has shown that the MT cytoskeleton in beta cells is actually arranged such that there is a central, non-directional meshwork with MTs predominantly perpendicular to the membrane, while there is a peripheral bundle of MTs lying parallel to the membrane (Zhu et al., 2015). Not only does this parallel bundle function to withdraw IGs away from sites of secretion (Bracey et al., 2020), but glucose stimulation promotes MT depolymerization (Zhu et al., 2015; Ho et al., 2020; Trogden et al., 2021) and MT depolymerization in turn promotes insulin secretion (Zhu et al., 2015). Moreover, we found that MT depolymerization specifically enhances clustered secretion at hot spots, including via increased number of secretion events per hot spot and activation of additional secretion hot spots, often in otherwise non-secreting beta cells (Trogden et al., 2021). We found that such MT-dependent activation of secretion hot spots is an essential factor contributing to the functional heterogeneity of beta cells (Benninger and Kravets, 2022). These findings raise the question of how MTs exert their negative regulatory role in directed insulin secretion.

In this study, we apply total internal reflection fluorescence (TIRF) microscopy and quantitative analysis of vascular ECM-associated insulin secretion patterns in intact mouse during first phase glucose-stimulated insulin secretion (GSIS) in correlation with ELKS and LL5β patterns and MT configuration at secretion sites. Our results indicate that active directed secretion is concentrated to regions at the margins of ELKS/LL5β patches which are also devoid of MTs in part due to MT depolymerization prior to secretion. We put forward a model where ELKS/LL5β-dependent MT anchoring promotes IG positioning to the proximity of secretion sites, but both anchoring patches and MTs interfere with secretion events themselves.

## Results

### ELKS patches display secretory heterogeneity

Since ELKS is considered a necessary molecular player involved in targeted secretion to the vasculature, we performed a detailed analysis of the spatial distribution of secretion events in relation to ELKS accumulation at the plasma membrane. We took advantage of an experimental model where whole-mount intact mouse islets with a GFP-ELKS knock-in (Figure S1A-C) are attached to the glass coated with the vascular ECM (Zhu et al., 2015; Ho et al., 2023a). Usage of vascular ECM enriched in laminin and collagen IV allows for our high confidence that we are recapitulating the *in vivo* interface of beta cells with endothelial-produced ECM (basement membrane), the only ECM type amply present within islets. In this experimental model, ELKS patches are assembled at the interface with the vascular ECM, similar to the patches at the capillaries within the islet (Figure S2A-D). This allows us to analyze a large area of ELKS at the vascular ECM interface by TIRF microscopy.

In line with previous groups’ observations (Ohara-Imaizumi et al., 2005; Low et al., 2014), ELKS at vasculature-cell interfaces formed distinct patches (Figure 1B). To confirm the specificity of these patches by their spatial relationships with integrin-dependent focal adhesions (Lansbergen et al., 2006; Bouchet et al., 2016, Fourriere et al., 2019; Noordstra et al., 2022), we show that ELKS localizes adjacent to actin fiber termini detected in transgenic Halo-Lifeact/knock-in GFP-ELKS mouse islets (Figure S3A-B). We further use NSPARC and TIRF microscopy to confirm that ELKS localizes to focal adhesion plaques (Figure S3C-F) detected by localization of adhesion-enriched actin-binding protein mCh-Utrophin (mCh-UtrCH) (Belkin and Burridge, 1995; Galkin et al., 2002).

**Figure 1.**
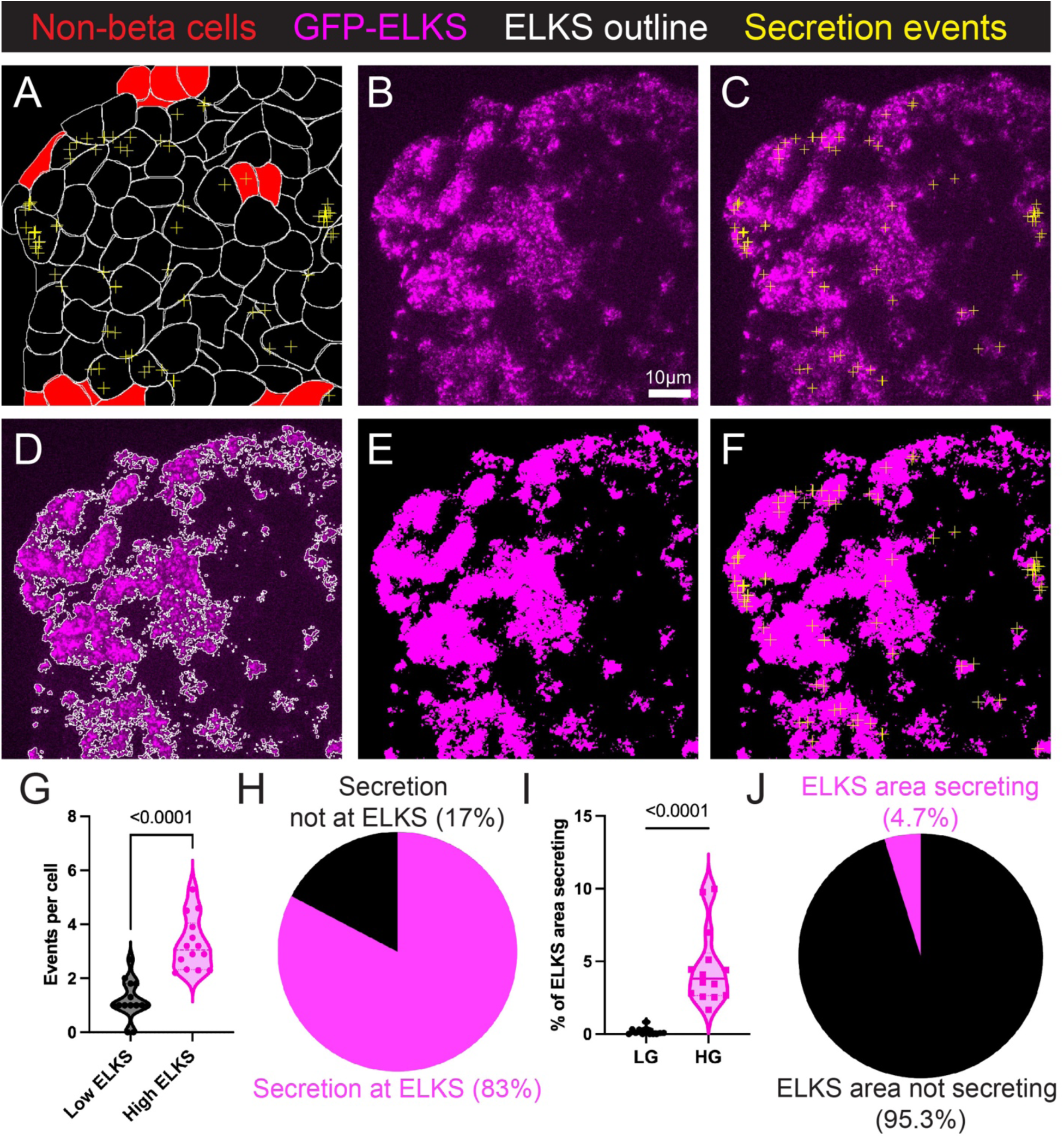
ELKS patches at the vascular ECM interface display secretory heterogeneity. A) Cell outlines at the ventral plane of an intact mouse islet (white) with insulin secretion events (yellow crosses), and non-beta cells (red). B) Single-plane TIRF image of knocked-in GFP-ELKS in the same islet (magenta). Scale bar = 10µm. See Video 1.2 for FluoZin-3 flashes. C) Secretion events (yellow crosses) overlaid on GFP-ELKS image, 8 minute movie (74 secretion events). D) Outline of GFP-ELKS patches (white). E) Thresholded ELKS patches (magenta). F) Thresholded ELKS patches (magenta) with secretion events (yellow crosses). G) Events per cell in low GFP-ELKS-expressing cells as compared to high GFP-ELKS-expressing cells. Student’s t test, P<0.05. N=14 high glucose (HG)-treated islets. H) Summarized data of secretion events occurring at GFP-ELKS-positive patches versus GFP-ELKS-negative beta cell area. N=14 HG islets. I) Percent of GFP-ELKS patch area (as in D-F) secreting in low glucose (LG) and HG. Student’s t test, P<0.05. N=12 LG, 14 HG islets. J) Summarized data of GFP-ELKS area secreting in HG. N=14 islets, n=33-226 secretion events per islet.

To analyze the secretory capacity of ELKS patches, we induced insulin secretion in the attached islets by high glucose. We capture real-time secretion events at this interface using the cell-impermeable Zn^2+^ binding dye, FluoZin-3, and imaging using TIRF microscopy to capture these events at this thin interface (Figure S4A-K, Video 1.1). In this technique, the center of mass of each FluoZin-3 flash is considered to indicate the location of the opening pore of the corresponding exocytic event. Coordinates of the secretion event are calculated with sub-pixel resolution by the same principle as super-resolution localization microscopy (Khater et al., 2020), following a Gaussian distribution of intensity (Figure S5A-O). Although several of these flashes are on the larger size (1-2µm), the majority have a radius of <1µm allowing for a high accuracy fitting of the flash center (Figure S4L-O) and high temporal localization accuracy in hundreds of milliseconds (Figure S4D-G, P). Given that the size of insulin granules is approximately 0.2-0.3µm (Fava et al., 2012), a potential error in the location of the opening pore on a scale of nanometers is negligible. Thereafter, we allocate the coordinates of the secretion events to the first frame of GFP-ELKS prior to FluoZin-3 treatment, as ELKS patch location and morphology is not changed detectably during the eight-minute acquisition period (Figure S6A-B; Noordstra et al., 2022). We can also define beta cells and non-beta cells using a beta-cell-specific nuclear probe and cell outlines using brightfield microscopy (Figure S2E-G’).

We have analyzed the distribution of insulin secretion events relative to ELKS localization in the TIRF plane (Figure 1C-F). Our analysis found that beta cells with high ELKS content secrete significantly more efficiently than those with low ELKS content (Figure 1G). Moreover, the majority of secretion events (83% vs 17%) occur at ELKS patches (Figure 1H) thereby confirming the importance of ELKS as a secretion-promoting factor as shown previously (Ohara-Imaizumi et al., 2005; Low et al., 2014). However, we made an unexpected finding that not all ELKS patches secrete (Video 1.2). In fact, high glucose only stimulated secretion within less than 5% of the total ELKS patch area (Figure 1I, J). In addition to this finding of ELKS heterogeneity, we observe that some cells secrete more than others, which is in line with our group’s and others’ previous findings that beta cells are highly heterogeneous in their secretion capacity (Van Schravendijk et al., 1992; Giordano et al., 1993; Wojtusciszyn et al., 2008; Trogden et al., 2021). As indicated here, however, the size of ELKS patches does not define preferred secretion per cell and does not alone underlie functional heterogeneity of beta cells.

Thus, our data are consistent with previous findings that ELKS is involved in targeted insulin secretion at the vascular interface, but at the same time, we show that ELKS patches display a surprising secretory heterogeneity: some ELKS patches secrete while most do not.

### LL5β patches display secretory heterogeneity

In attempting to determine whether other markers of directed secretion might be better predictors of secretion sites, or if they, too, experience secretory heterogeneity, we next turned to the ELKS binding partner LL5β (Lansbergen et al., 2006), which is required for clustering of insulin granule docking complexes (Noordstra et al., 2022) and thus making it a plausible candidate for influencing directed secretion in beta cells.

We used the same techniques as in Figure 1 to identify beta cells and secretion events in intact mouse islets transduced with lentivirally-expressed RFP-LL5β (Figure 2A-I, Video 2). Like ELKS, RFP-LL5β patches display very little dynamics over the course of acquisition (Figure S6C-D). Additionally, due to this technique, RFP-LL5β is expressed mosaically, in selected beta cells (Figure S1D-G). The importance of LL5β in the secretion process is confirmed by our observation that cells ectopically expressing RFP-LL5β show slightly higher secretion than cells without RFP-LL5β expression (Figure 2J). However, we cannot distinguish whether secretion events occur at endogenous LL5β patches or not. Accordingly, we exclude RFP-LL5β-negative cells from the analysis of secretion events positioning relative to RFP-LL5β patches.

**Figure 2.**
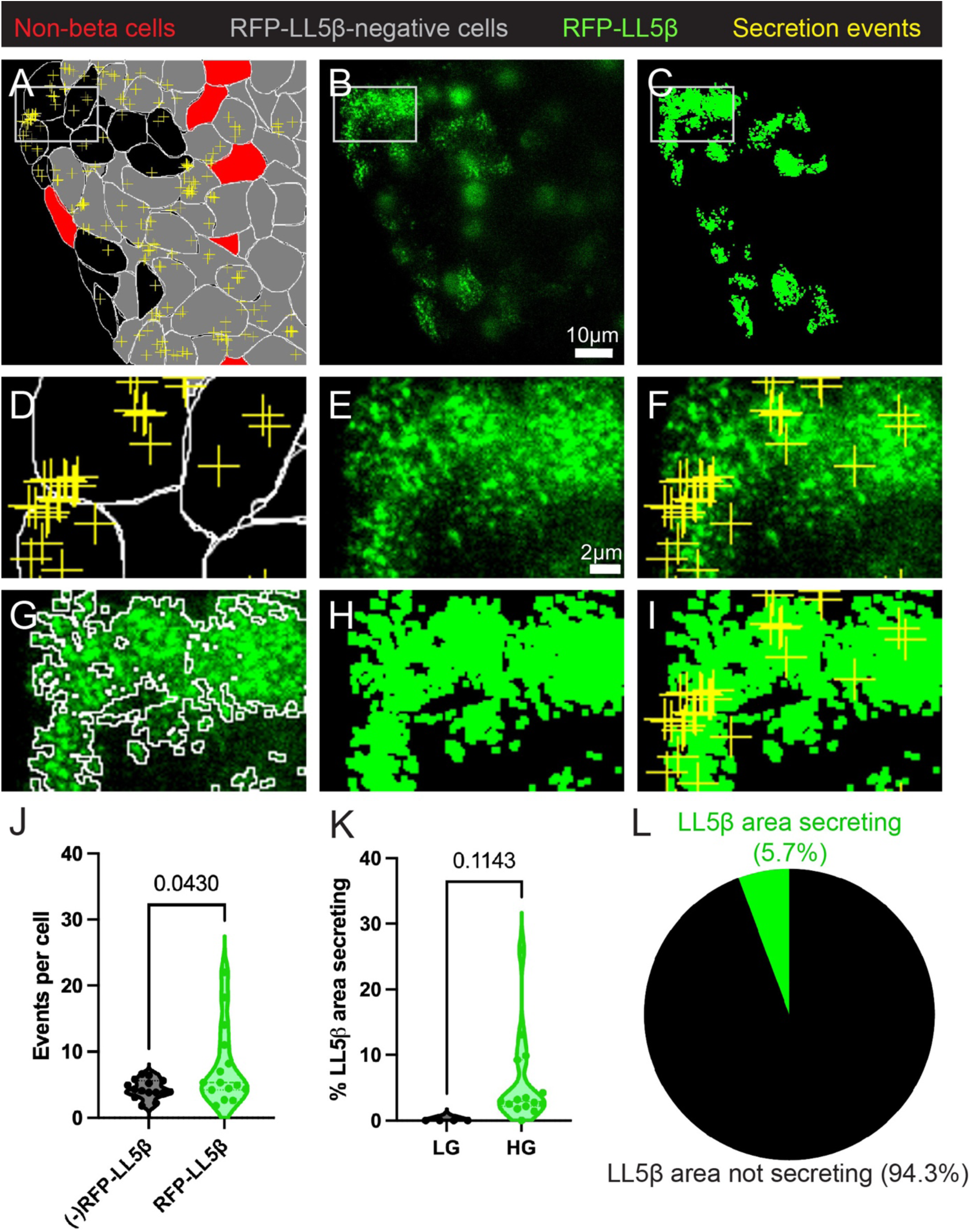
LL5β patches at the vascular ECM interface display secretory heterogeneity. A) Islet cell outlines (white) with insulin secretion events (yellow crosses), and non-beta cells (red). Non-analyzed cells, gray. From an 8 minute movie (82 secretion events) B) Single-plane TIRF image of ectopic RFP-LL5β in the same islet (green). Scale bar = 10µm. C) Thresholded RFP-LL5β patches (green). D-I) Insets of A-C. D) Islet cell outlines (white) with insulin secretion events (yellow crosses); inset cells express RFP-LL5β. E) RFP-LL5β (green). Scale bar = 2 µm. F) RFP-LL5β (green) and secretion events (yellow crosses). See Video 2 for FluoZin-3 flashes. G) Outline of RFP-LL5β patches (white). H) Thresholded RFP-LL5β patches (green). I) Thresholded LL5β patches (green) with secretion events (yellow crosses). J) Events per cell in cells not expressing RFP-LL5β as compared to those expressing RFP-LL5β. K) Percent of RFP-LL5β patch area (as in G-I) secreting in LG and HG. Student’s t test, P<0.05. N=9 LG, 15 HG islets. L) Summarized data of RFP-LL5β area secreting in HG. N=15 islets, n=49-599 secretion events per islet.

In cells with ectopic RFP-LL5β expression, we observe multiple secretion events at LL5β patches (Figure 2D-I, insets). However, less than 6% of total LL5β patch area secretes in response to high glucose stimulus (Figure 2K-L) thus indicating secretory heterogeneity very similar to ELKS patches. We further corroborate these results in cells co-expressing GFP-ELKS and RFP-LL5β. As expected, these two proteins often co-localize in the same cortical patches which show the same secretory heterogeneity as patches detected by each protein separately (Figure S7A-L). This indicates that while both proteins serve as components of cortical secretory patches, LL5β does not serve as a defining factor for ELKS patches’ capacity to secrete.

### Secretion events preferentially occur at margins of ELKS patches

After excluding LL5β as a factor promoting secretion within ELKS patches, we proceeded to analyze the characteristics of ELKS patches themselves that might define their heterogeneity. In particular, we addressed whether a higher fluorescent pixel intensity of GFP-ELKS (a proxy for the concentration of ELKS molecules) correlates with a higher probability of secretion (workflow, Figure 3A). As expected, secretory events predominantly occur at ELKS-positive pixels, which would be any value above 0 in the whole-cell measurements (Figure 3B). However, the comparison of the ELKS intensity at the secretion points with ELKS intensity frequency within patches showed a slight shift of secretion toward lower ELKS intensity within patches (Figure 3C). In particular, we find multiple secretion events in high-intensity regions (>0.2 intensity, Figure 3D-D’, Movie 3.1), a large percentage of secretion events in the regions of low-intensity ELKS (0.1-0.2 intensity, Figure 3E-E’, Movie 3.2) and a fraction of events (∼15%) at background intensity (0 intensity, Figure 3F-F’, Movie 3.3).

**Figure 3.**
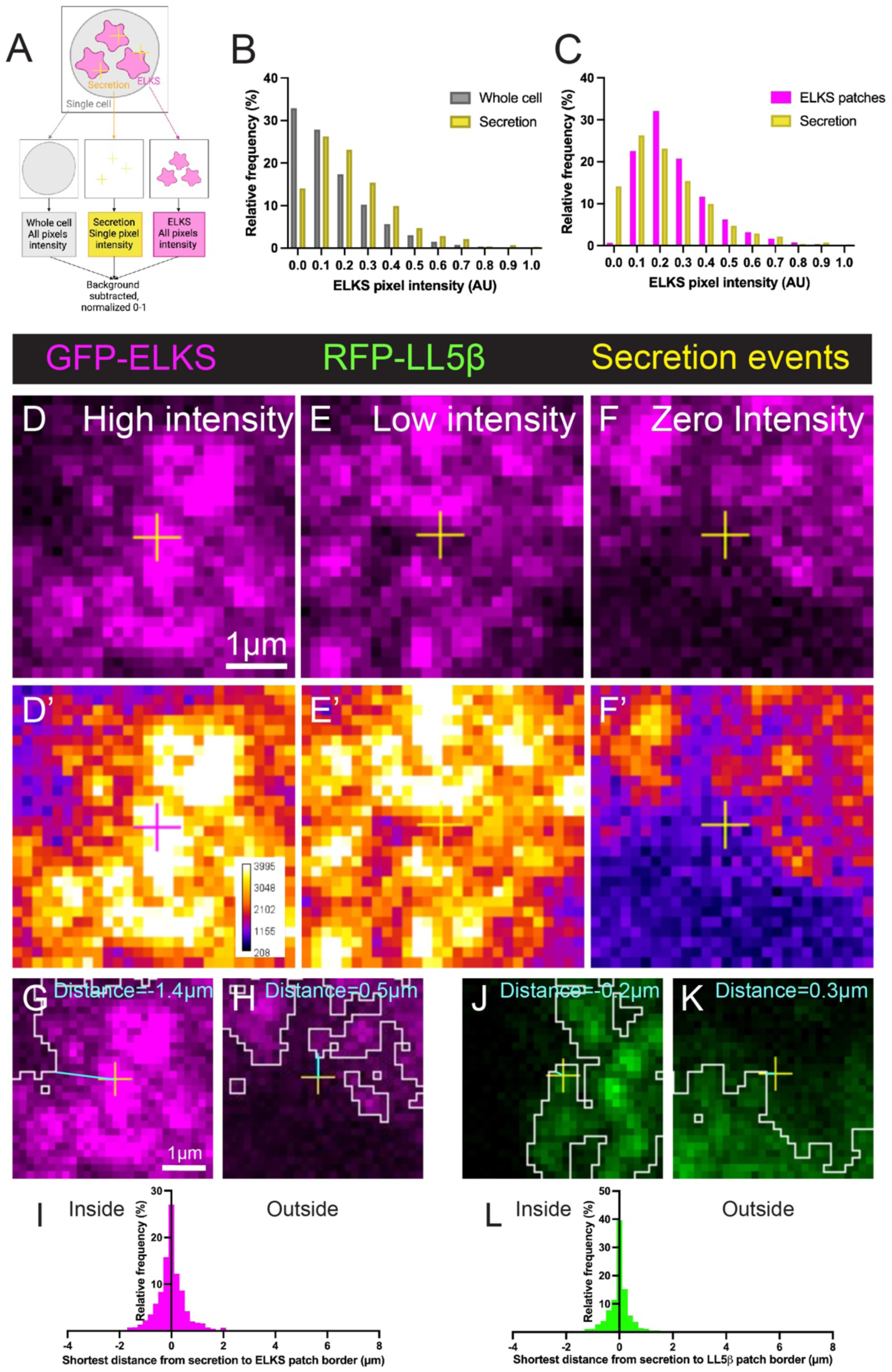
Secretion events preferentially occur at marginal regions of ELKS patches. A) Schematic showing the workflow of splitting cell components and taking all pixel intensity of ELKS (magenta), single pixel intensity of secretion events (yellow), and whole cell pixel intensity (gray). B) Percent relative frequency of normalized pixel intensity for all pixels in a cell (gray) and the pixel at the point of secretion (yellow). C) Percent relative frequency of normalized pixel intensity for all pixels within ELKS patches (magenta) and the pixel at the point of secretion (yellow). N=14 islets, n=33-226 secretion events per islet; 1,135,338 whole cell pixels, 523,416 ELKS pixels, 1,484 secretion event pixels. D) Crop of a TIRF image of a whole mouse islet expressing knocked-in GFP-ELKS (magenta). Secretion event (yellow cross) occurs at a region of high-intensity ELKS. Scale bar = 1µm. E) Secretion event (yellow cross) occurs at a region of low-intensity ELKS, at the margin of an ELKS puncta. F) Secretion event (yellow cross) occurs at a region of background intensity, still near an ELKS patch. D’-F’) Heat maps of ELKS expression corresponding to D-F). See Videos 3.1-3.3 for FluoZin-3 flashes. Secretion event in magenta for D) for contrast. G) Shortest distance between internal secretion event and ELKS patch edge, 1.4µm. H) Shortest distance between external secretion event and ELKS patch edge, 0.5µm. I) Histogram of shortest distances between secretion events internal and external to ELKS patches. Inside patch = negative, outside patch = positive. J-K) Shortest distance examples between secretion events and LL5β patches edges. L) Histogram of shortest distances between secretion events internal and external to LL5β patches.

A closer visual analysis of secretion event location suggested that the low-intensity ELKS regions where secretion occurred were located at the edges or adjacent to ELKS patches. To more thoroughly explore the spatial relationship between secretion events and ELKS patches, we further performed a shortest distance analysis between events and the edges of ELKS patches as defined by thresholding. We found that ∼72% of events occur within 0.3µm, and ∼27% of events within 0.1 µm of an ELKS patch edge (Figure 3G-I). The events were found equally either inside or outside the patch (for quantification purposes, locations inside the patch were assigned negative, and outside the patch assigned positive values). A much smaller percentage occur more central to the ELKS patch or outside, consistent with our pixel intensity analysis. We showed the same pattern occurs for LL5β (Figure 3J-L) with ∼80% of events occurring within 0.3µm and ∼40% of events occurring in this short -0.1 to 0.1 µm range of an LL5β patch. These data indicate that secretion events preferentially occur at the margins of ELKS patches where ELKS/LL5β concentration is modest. These data lead to the hypothesis that while high-content ELKS accumulations may play a role in preparation for secretion, for example, by promoting IG delivery to secretion sites or activation of calcium channels, the margins of ELKS patches provide the environment beneficial for exocytosis itself. In light of our confirmation here that ELKS patches assemble outside of actin bundle termini and focal adhesion plaques (Figure S3), our data support a model in which secretion location is indirectly defined by focal adhesion through positioning of ELKS patches and their margins.

### MT plus ends marked by KIF5B and capable of trafficking IGs do not associate with insulin secretion event sites

We next turned to explore ELKS/LL5β-dependent MT regulation as a potential factor in secretory heterogeneity of patches. Our group has already shown that MT depolymerization activates additional secretion hot spots (Trogden et al., 2021). This, along with the finding that LL5β can anchor MT plus ends at the cortex (Lansbergen et al., 2006; Basu et al., 2015) suggested to us that MT plus ends may play a regulatory role in determining whether or not these patches actively secrete.

The traditional method of visualizing MT plus ends by fluorescently-labeled end-binding (EB) protein comets (Akhmanova and Hoogenraad, 2005; Sanchini et al., 2023) presents a unique challenge in primary beta cells due to their high MT stability (Zhu et al., 2015), such that most MT ends are not polymerizing and thus are not detected by EB comets. In light of this challenge, we use an approach where molecular motor tracking is used to label MT polarity and molecular motor dwelling/accumulation is used to track MT end location (Noda et al., 2001; Guardia et al., 2016; Tas et al., 2017). Specifically, we tracked the movement of KIF5B motor domain in the TIRF optical plane to detect plus ends of MTs which are suitable for Kinesin-1 (KIF5B)-mediated transport of IGs. For single-molecule tracking of KIF5B motors, we utilized the existing SunTag system, which allows high numbers of fluorophores to bind each KIF5B motor (Tanenbaum et al., 2014). This allows us to observe the directionality of KIF5B motor movement of 47% of KIF5B tracks (“directed” motion high-displacement tracks, Figure 4A-C, Track 1, Video 4). 53% of KIF5B tracks presented low displacement and were interpreted as motor dwelling points at MT plus ends (KIF5B low-displacement tracks, Figure 4A-C, Track 2, Video 4). There was no significant difference in track displacement length between cells that secrete and cells that do not secrete within the same islet (Figure 4C). Using the last track point of both track groups to proxy the MT plus ends, we also analyze the distances between MT plus ends and secretion events detected by FluoZin-3 secretion assay (Figure 4D-G). Our quantification indicates that less than 1% of secretion events take place within 0-1µm of KIF5B track endpoints while the majority of secretion occurs within 1-6µm of those endpoints (Figure 4H). A similar relationship was found between secretion hot spots and clusters of MT plus ends at the cell periphery (Figure 4I-J). These data together suggest that secretion is not immediately associated with either single MT plus ends or MT plus end accumulations at the beta cell periphery thus raising further questions about the relationship between MT plus-end-directed delivery of IGs and secretion event location.

**Figure 4.**
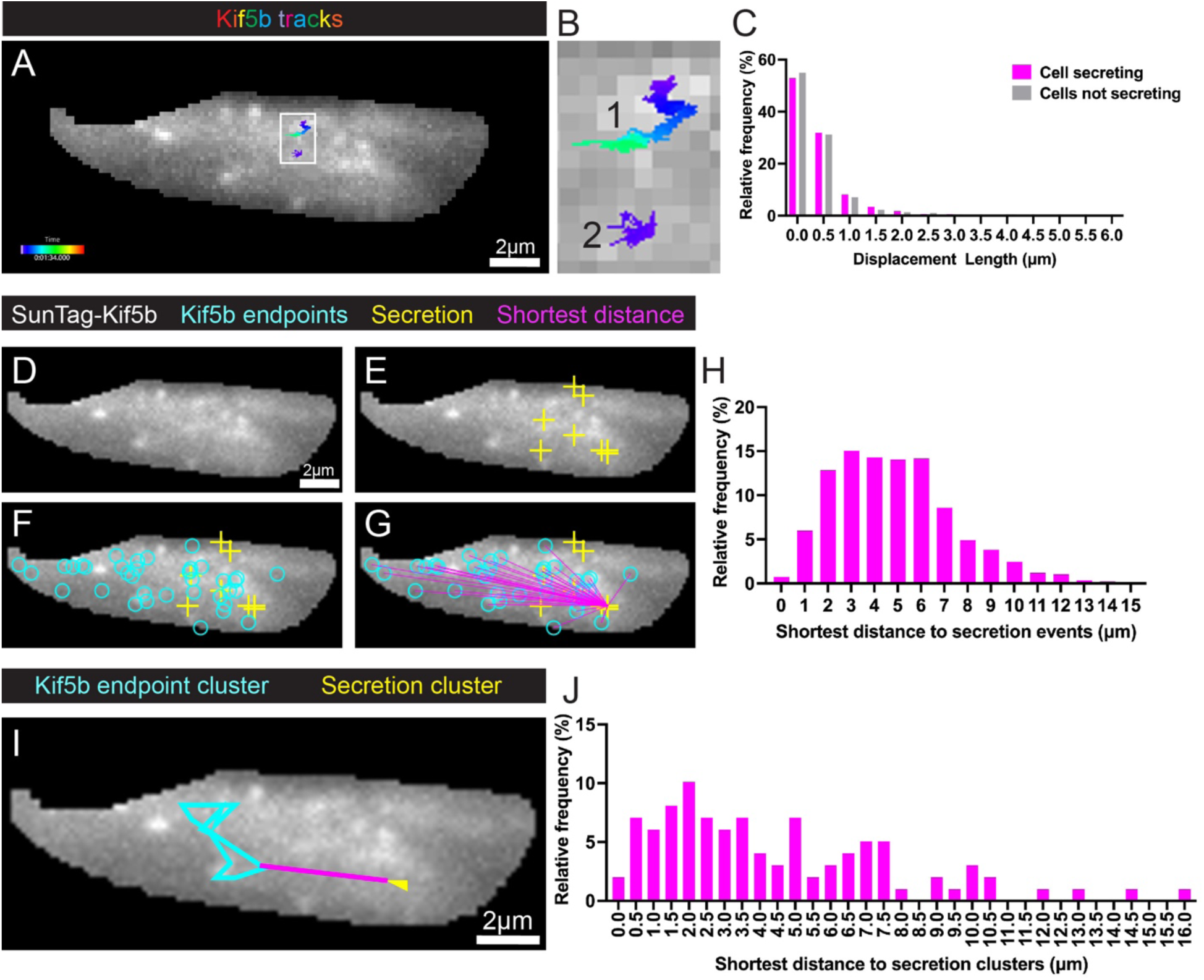
MT plus ends marked by SunTag-KIF5B and capable of trafficking IGs do not associate with insulin secretion event sites. A) Example of SunTag-KIF5B motor tracks in a masked-out beta cell within an islet, TIRF microscopy. Corresponds to Video 4. Scale bar = 2µm. B) Inset of A) with a long-displacement length track (1) and a short-displacement length track (2). C) Percent relative frequency of displacement lengths of tracks compared between secreting cells and non-secreting cells. Not significant, P<0.05, student’s t test. N=14 islets, n=2,439 displacement lengths. D-G) Shortest distance analysis. Scale bar = 2µm. D) SunTag-KIF5B (gray). E) Insulin secretion events (yellow crosses) identified by FluoZin-3, in addition to D). F) KIF5B track endpoints (cyan circles) in addition to E). G) Example shortest distance between an insulin secretion event and all KIF5B track endpoints in a cell (magenta), in addition to F). H) Percent relative frequency of shortest distances between secretion events and KIF5B track endpoints. N=14 islets, n=6,429 secretion events. I) Example of a shortest distance measurement (magenta) between a cluster of KIF5B track endpoints, represented as a polygon (cyan) and a secretion cluster polygon (yellow). J) Percent relative frequency of shortest distance between KIF5B track endpoint clusters and secretion clusters. N=14 islets, n=98 secretion clusters.

### MT minus ends are not associated with insulin secretion event sites

After determining that MT plus ends are not associated with secretion events, we sought to understand whether MT minus ends have any association. We and others have previously observed that MT minus ends in pancreatic beta cells can be found at the cell periphery (Müller et al., 2021; Ho et al., 2023b). Now, using expression of MT minus-end-binding protein GFP-CAMSAP2 as a marker, we not only confirm these observations (Figure 5A-B), but also find multiple MT minus ends in the TIRF microscopy optical plane where beta cells are in contact with the vascular ECM (Figure 5C-F). Thus, it is possible that minus end-directed motors could carry IGs to directed secretion sites at the beta cell periphery via cytoplasmic dynein (Varadi et al., 2003) or minus-end-directed kinesin motors. In the interest of exploring whether MT minus ends have any relation to secretion events, we analyzed the distances between GFP-CAMSAP2 and FluoZin-3 flashes. We show by comparing the secretion events with minus ends (Figure 5C-F, Figure S8, Video 5) that there is no significant pattern at which MT minus ends are positioned relative to secretion events (Figure 5G). We conclude, based on both the plus and minus end data (Figs. 4 and 5), that MT ends do not define the precise location of secretion events and therefore cannot underlie the secretory heterogeneity within ELKS/LL5β patches (Figs. 1-3).

**Figure 5.**
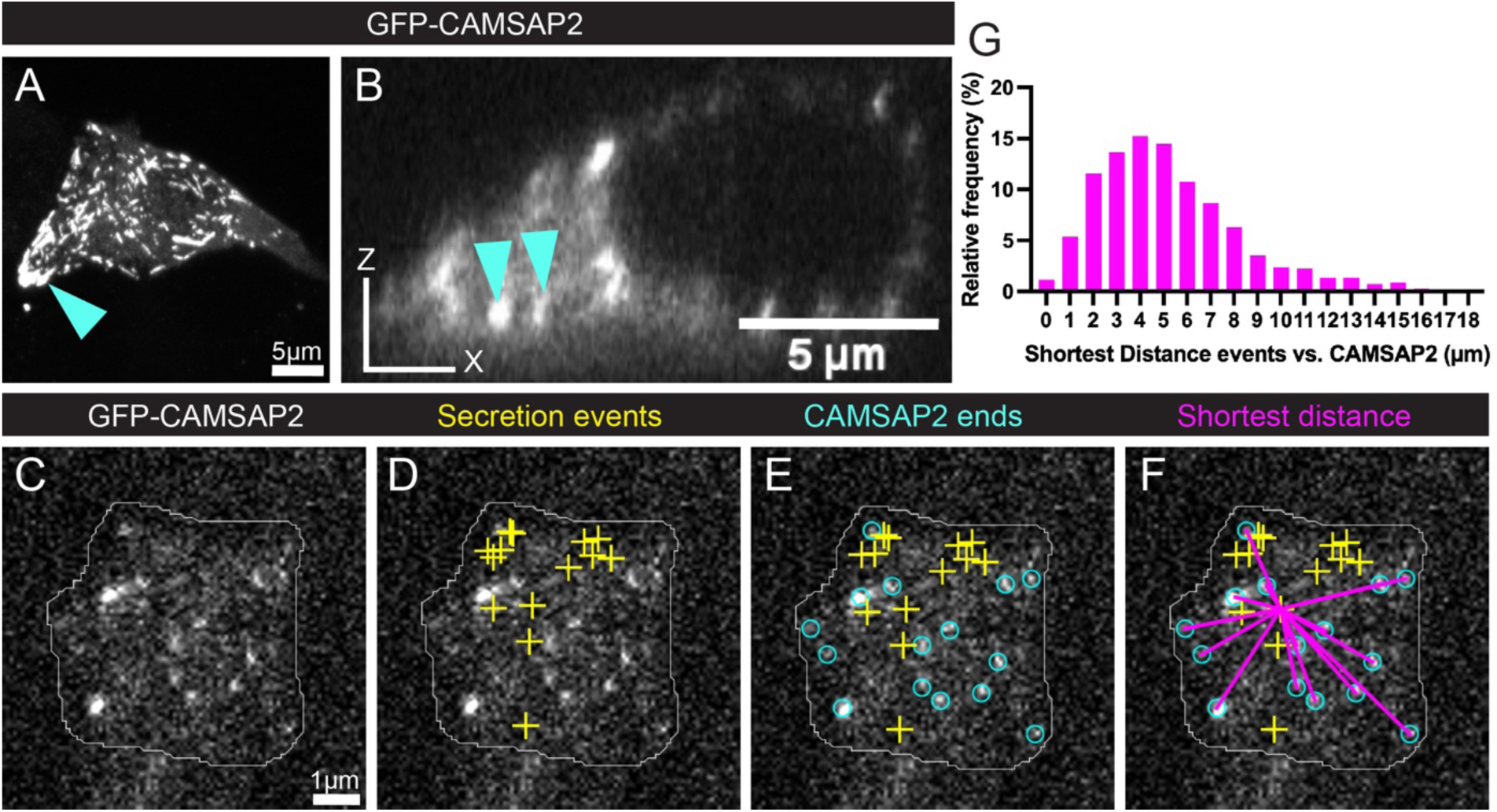
MT minus ends marked by CAMSAP2 are not associated with insulin secretion event sites. A) MIN6 beta cell maximum intensity projection, expressing ectopic GFP-CAMSAP2, in which many CAMSAP2 stretches and puncta localize to the periphery (cyan arrowhead). Scale bar = 5µm. B) MIN6 beta cell, single slice XZ (side) view, showing CAMSAP2 at the bottom of the cell (cyan arrowheads). Laser scanning confocal microscopy. C-F) Individual islet beta cell expressing ectopic GFP-CAMSAP2 imaged with TIRF microscopy, exemplifying CAMSAP2 puncta marking MT minus ends. C) cell (outlined in white) expressing CAMSAP2 (white). Scale bar = 2µm. D) insulin secretion events (yellow crosses) identified by FluoZin-3, in addition to C). E) MT minus ends (cyan circles) in addition to D). F) example shortest distance between an insulin secretion event and all MT minus ends in a cell (magenta), in addition to E). G) Histogram of the percent relative frequency of shortest distance between insulin secretion events and CAMSAP2 puncta, showing that only 1.13% of events fall between 0-1µm of MT minus ends. See Video 5 for FluoZin-3 flashes. n=1955 shortest distance measurements.

### The majority of secretion events occur in regions devoid of MTs

Having determined that MT end location does not define the secretory capacity of ELKS and LL5β patches, we turned to evaluate whether other configurations of MTs are specifically associated with secretion sites. Our lab has previously shown that peripheral MT arrays oriented parallel to the plasma membrane serve to traffic IGs away from sites of secretion and that glucose-induced destabilization of those arrays promotes secretion (Zhu et al., 2015; Ho et al., 2020; Trogden et al., 2021). Accordingly, we proceed to test whether MT lattices parallel to the plasma membrane influence secretion at a single secretion event level as compared to MT ends and areas devoid of MTs. Using NSPARC super-resolution microscopy, we concurrently visualize Halo-tubulin, GFP-ELKS, and RFP-LL5β in an islet beta cell and show three existing configurations of MTs relative to these puncta: lattice-bound, end-on, and not associated (N/A) (Figure 6A-A”). We also demonstrate our ability to visualize punctate or short stretches of MTs at the bottom of the cell as compared to the whole cell in NSPARC (Figure 6B-D).

**Figure 6.**
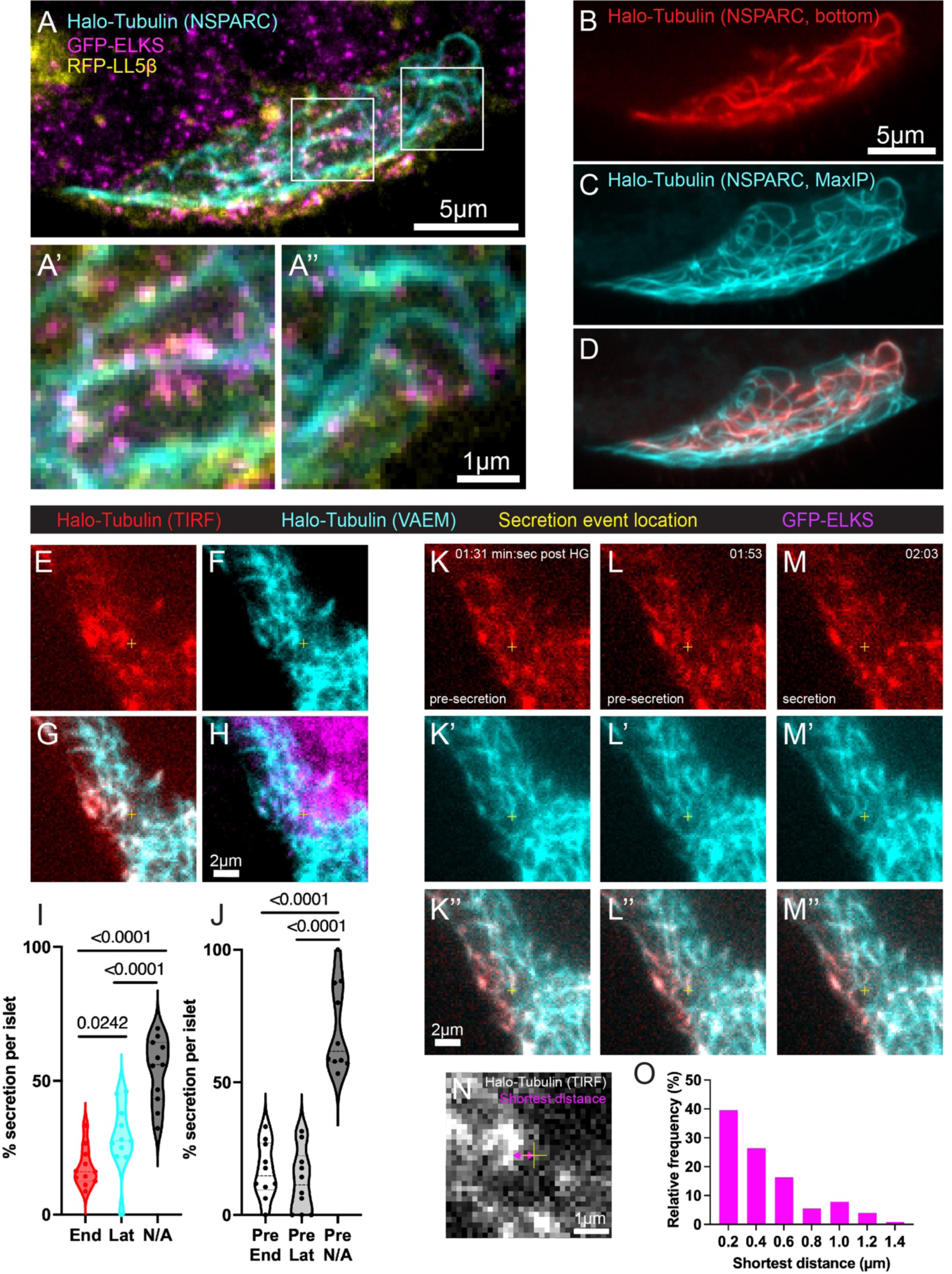
The majority of secretion events occur in regions devoid of MTs. A) Islet beta cell expressing ectopic Halo-Tubulin, ectopic RFP-LL5β, and knocked-in GFP-ELKS, imaged in super-resolution using NSPARC to highlight A’-A”) lattice-bound, N/A, end-on configurations. B) Cell bottom showing fragments of Halo-tubulin-labeled MTs and C) MaxIP to show full MT configuration. D) Composite of B) and C). E-H) Example of N/A configuration analyzed by TIRF/VAEM imaging in combination with the FluoZin-3 assay. Yellow cross indicates secretion location for all panels. E) Halo-Tubulin in the TIRF Plane (red). F) Halo-Tubulin in the VAEM plane (cyan). G) TIRF and VAEM planes acquired concurrently during imaging, in addition to H) GFP-ELKS (magenta) to confirm presence of secreting patches. I) Percent of secretion per islet occurring in an end-on (End), lattice-bound (Lat), or non-associated (N/A) configuration relative to local MTs. Student’s t test, P<0.05. N=11 islets, n=14-129 secretion events. J) O) Percent of secretion per islet occurring in either an end-on, lattice-bound, or non-associated configuration relative to MTs (Pre End, Pre Lat, and Pre N/A, respectively), prior to an N/A secretion event. Student’s t test, P<0.05. N=11 islets, n=14-129 secretion events. K-M”) Montage of a MT moving away from a secretion site in an islet beta cell, imaged in TIRF and VAEM. K-M) TIRF plane. K’-M’) VAEM plane. K”-M”) Composite. Example shows a MT overlapping a future point of secretion at time 01:31 min:sec in K-K”), beginning to move away from the future point of secretion at time 01:53 in L-L”), and finally absent at the point of secretion at time 02:03 in M-M”). Corresponds to Video 6. P) Example of a shortest distance measurement (magenta arrows) between secretion and MT in TIRF. Q) Percent relative frequency of shortest distances between secretion events and their nearest MTs. Scale bar = 1µm. Student’s t test, P<0.05. N=11 islets, n=129 secretion events.

To assess MT organization at the vascular ECM attachment sites, we visualize MTs labeled with Halo-Tubulin by TIRF microscopy coupled with variable angle epifluorescence microscopy (VAEM). TIRF imaging allows for visualization of short MT stretches closely associated with the plasma membrane. VAEM, also known as near-TIRF, provided a deeper penetration of the excitation light than TIRF and allows for imaging of MT networks within the thicker optical slices above the cell membrane (Konopka and Bednarek, 2008). The combination of two techniques allows us to determine whether MT stretches detected in TIRF correspond to specific locations within long MTs detected by VAEM: either MT ends or regions of sub-membrane MT lattices (Figure S9).

We combine this approach with our FluoZin-3 secretion assay to determine whether secretion events are associated with a specific MT configuration (Figure 6E-H, Figure S10). We classify MT configurations similarly as above, but specifically at secretion event locations into three groups: MT ends at the plasma membrane (end-on), lattices of MTs at the plasma membrane (lattice-bound), and no MTs at the plasma membrane (No association [N/A]). Similar configurations can be found at ELKS patches in TIRF (Figure 6H, Figure S10A-I).

We find that the majority of all secretion events (average 54% per islet) occur in regions locally devoid of MTs (in other words, in regions displaying an N/A configuration, Figure 6I). This is consistent with our finding (Figs. 4 and 5) that MT ends are not specifically associated with sites of secretion. For cases with the N/A MT configuration at the moment of secretion, we also analyzed MT configuration prior to secretion. We found that MTs displayed end-on (17%) or lattice-bound (13%) configuration before being removed (Figure 6J). In 70% of these cases, MTs were not found at the site prior to secretion for the duration of high glucose stimulation. MT removal in these cases could be a result of MT depolymerization (Figure S10J-L”) or whole MT relocation (MT sliding) (Figure 6K-M”, Video 6.1-6.2). In either scenario, it is possible that IGs are being delivered to the secretion sites along MTs, which then depolymerize or slide away to allow for secretion to occur. This suggests that IGs may be delivered to those sites before the glucose-induced secretion triggering, consistent with our previous work showing that MT depolymerization promotes insulin secretion. Interestingly, we find (Figure 6N) that 40% of N/A secretion events occur relatively close (within 0.2-0.4µm) of the nearest MT (Figure 6O), suggesting that MT removal even to a short distance might be sufficient to allow for secretion.

## Discussion

In this paper, we present our analyses of factors essential for the first phase of directed GSIS toward vascular ECM at the high/super-resolution level. While confirming other groups’ findings that insulin secretion occurs at cell domains enriched with ELKS and LL5β (Ohara-Imaizumi et al., 2005; Ohara-Imaizumi et al., 2019b; Noordstra et al., 2022), we report secretory heterogeneity within ELKS/LL5β patches, indicating that the presence of these factors is not sufficient to define secretion hot spots. Surprisingly, we find that secretion preferentially occurs at the margin of an ELKS/LL5β patch, often at areas of lower ELKS content or between brighter ELKS puncta. This finding provides precise specificity to the secretion event location, while previously secretion was assumed to occur randomly at all ELKS accumulations at the focal adhesion periphery (Noordstra et al., 2022). Also, in agreement with our previous finding that MT depolymerization enhances phase 1 of GSIS (Trogden et al., 2021), we report that the absence of MTs from the sub-membrane region is a good predictor for local secretory activity. We further report that MT ends and therefore IG delivery sites are distinct from final sites of secretion. These findings significantly expand our understanding of how insulin secretion is spatially regulated at the sub-cellular level. We propose a model for site-specific regulation of insulin secretion based on our new data combined with the latest findings in the field (Figure 7).

**Figure 7.**
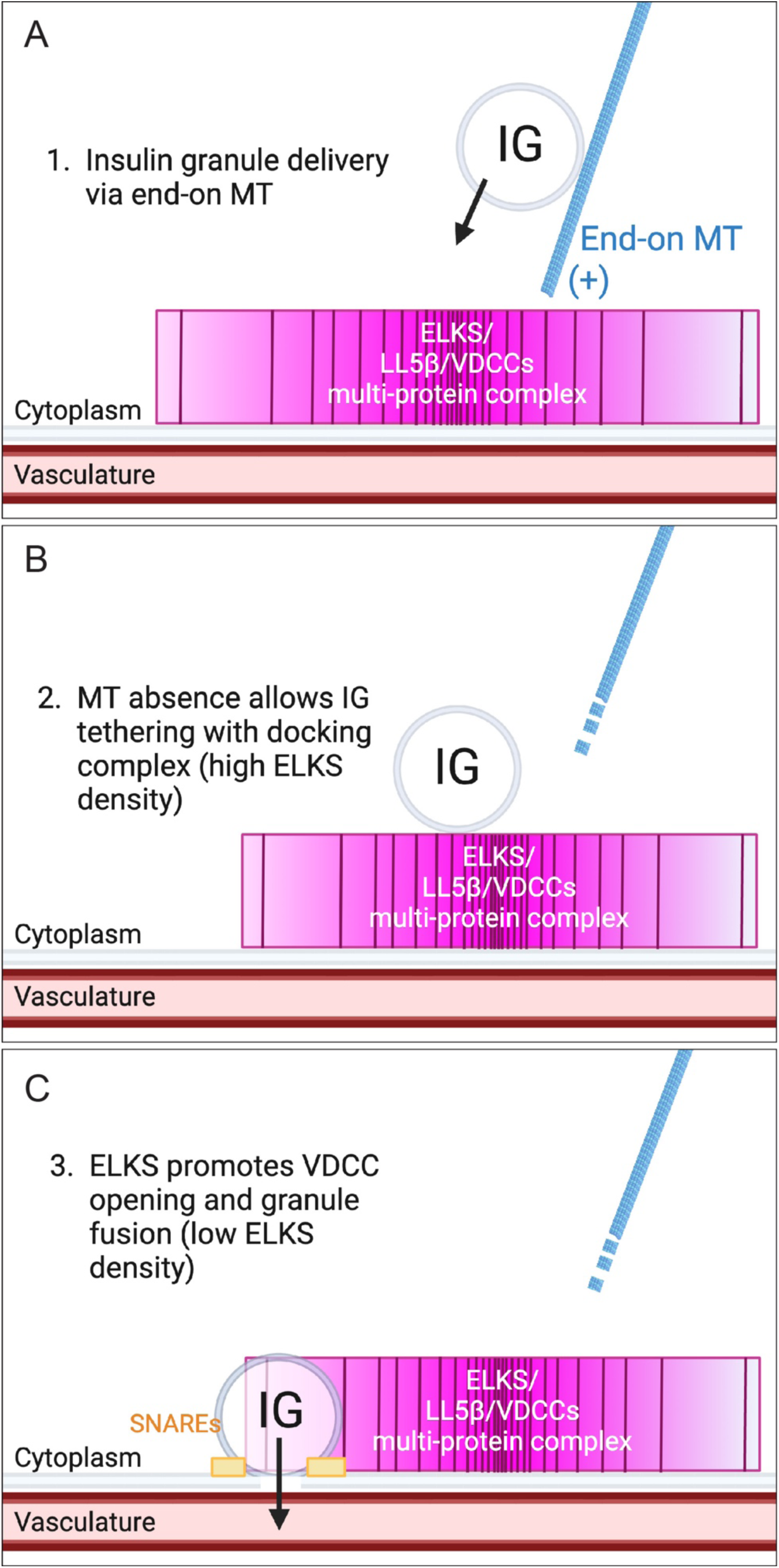
Regions devoid of MTs and at the margins of ELKS patches are strong predictors of secretion sites. A) Schematic detailing IG delivery to site of secretion via a MT bound to the ELKS/LL5β multi-protein complex (including VDCCs). Gradient and positioning of magenta multi-protein complex indicates intensity of ELKS signal and thereby ELKS concentration. B) MT absence, whether via depolymerization, sliding, or fragmentation, allows IG tethering at the multi-protein complex. C) ELKS promotes opening of VDCCs, allowing for granule fusion at the edge of ELKS regions.

Our novel finding that ELKS patches have secretory heterogeneity is interesting in the context of existing literature, that has suggested that all ELKS patches promote secretion (Ohara-Imaizumi et al., 2005; Low et al., 2014; Ohara-Imaizumi et al., 2019b). We found not only that a large percentage of the ELKS patch area does not secrete, but that secretion events rarely localize to the central parts of ELKS accumulations. One potential explanation is that ELKS function in secretion is associated with a rapid molecular exchange of ELKS protein within patches, facilitated by high glucose stimulation (Noordstra et al., 2022). Because ELKS molecules cluster in a non-liquid-liquid phase separation manner (Noordstra et al., 2022), it is possible that molecular turnover is inefficient in the middle of high ELKS-content patches. Another possibility is that molecular complexes within ELKS patches interfere with secretion for steric reasons. ELKS, being a large scaffolding protein, binds many other proteins to create large complexes (Ohara-Imaizumi et al., 2005; Grigoriev et al., 2011; Sudhof, 2012) and is also known to physically separate presynaptic vesicle capture and exocytosis (Nyitrai et al., 2020). The exact diameter of this complex is unknown and difficult to characterize given its complexity, but given the number of large proteins involved it is likely that they are as large as or larger than the diffraction limit. This implies, using our sub-pixel resolution from the FluoZin-3 assay, that we may be observing secretion occurring just adjacent to ELKS hubs and the gaps in space are due to other associated proteins. This further fits in with previous findings that a few ELKS molecules will associate within a patch (Noordstra et al., 2022), and each likely binds tens of proteins. In addition, there may be more yet unknown machinery adjacent to the brightest patches of ELKS that could play a role in directed secretion. In light of prior findings which show the enrichment of docked IGs at the high intensity ELKS patches (Ohara-Imaizumi et al., 2005; Low et al., 2014; Ohara-Imaizumi et al., 2019b), such specific selection of the secretion location might have a high functional significance in defining which IGs out of the docked IG population have a higher probability to secrete.

In any of the scenarios proposed here, targeted secretion at sites adjacent to high-concentration ELKS patches must include delivery of IGs to these sites. Many studies have put forth the hypothesis that plus-end-directed trafficking along MTs, via Kinsein-1 (KIF5B), is the primary mode of transport for IGs from the Golgi or reserve pool to the readily releasable pool, docking sites, or sites of secretion (Varadi et al., 2003, Cui et al., 2011). Moreover, the MT plus end anchoring protein LL5β is needed for secretion (Noordstra et al., 2022) and its over-expression enhances the secretory capacity of individual beta cells (Figure 2). However, despite the prediction that plus end-mediated delivery of IGs would dictate the secretion sites and therefore ELKS/LL5β patch activity, we found that MT end location in the time frame of glucose stimulation does not correspond to the sites of secretion. In contrast, we discovered that most secretion events occur at regions of patches that are devoid of MTs. There are two potential non-exclusive explanations for this finding. The first explanation is based on our finding that secretion can be observed shortly following MT disappearance. Such disappearance can be explained by either MT depolymerization (Zhu et al., 2015; Ho et al., 2020), sliding away from the secretion site (Bracey et al., 2023), or MT fragmentation by severing enzymes such as katanin (proposed in Müller et al., 2021). Since our lab has shown that MTs have a net negative regulatory effect on insulin secretion by trafficking granules away from sites of secretion (Bracey et al., 2020), the second explanation is that IGs are delivered via MT-dependent delivery to the secretion sites by short-range actin-dependent trafficking via unconventional myosin (Ivarsson et al., 2005; Varadi et al., 2005; Arous et al., 2013; Tokuo et al., 2021).

Overall, we have found several important features of directed insulin secretion that were previously uncharacterized. Importantly, this study confirms that ELKS/LL5β complexes are important for secretion and is consistent with our lab’s previous findings that MT depolymerization locally promotes secretion. At the same time, our findings call for further exploration of mechanisms defining the location and efficiency of directed insulin secretion. Additionally, our study model holds certain limitations inherent to these methods and techniques that warrant addressing. In particular, we exclusively characterize first phase insulin secretion. It is challenging to assess the second phase of GSIS with our experimental model due to accumulating fluorescent FluoZin-3 background with growing Zn^2+^ concentration in the medium. However, the role that MTs play at ELKS during second phase secretion, specifically whether or not they continue to be absent from ELKS patches to allow for secretion, is of great interest.

## Materials & Methods

### Key resources table

| Reagent type (species) or resource | Designation | Source or reference | Identifiers | Additional information |
| --- | --- | --- | --- | --- |
| Genetic reagent (*Mus musculus*) | Ins-Apl mice | Stancill et al., 2019 |  | Histone 2B-mApple knocked into the *Ins2* locus |
| Genetic reagent (*Mus musculus*) | Ins-Apl/GFP-ELKS mice | Noordstra et al., 2022 |  | Homozygous Ins-Apl mice crossed with GFP-ELKS knock in mice |
| Genetic reagent (*Mus musculus*) | Ins-Apl/Halo-Lifeact/GFP-ELKS mice | Noordstra et al., 2022; VGER/Matthew Tyska, Vanderbilt University |  | Homozygous Ins-Apl/GFP-ELKS mice crossed with Halo-Lifeact transgenic mice |
| Chemical compound, drug | FluoZin-3, Tetrapotassium Salt, cell impermeant | Thermo Fisher Scientific, | Cat#: F24194 | Final concentration (20 μM) |
| Peptide, recombinant protein | Human ECM | Corning | Cat#: 354237 | Placenta-derived vascular ECM |
| Antibody | Rabbit Polyclonal anti-PHLDB2 | Invitrogen | Cat#: PA5-57176 | Antibody against LL5β |
| Antibody | Rabbit Polyclonal anti-ERC1 | Atlas Antibodies | Cat#: HPA019513 | Antibody against ELKS |
| Antibody | Mouse Monoclonal anti-GAPDH 6C5 | Santa Cruz Biotechnology | Cat#: sc-32233 |  |
| Antibody | IRDye 680 Goat anti-Rabbit | LI-COR BioSciences | Cat#: 926-68071 |  |
| Antibody | IRDye 800 Goat anti-Mouse | LI-COR BioSciences | Cat#: 926-32210 |  |
| Antibody | Goat anti-Rabbit 488 | Invitrogen | Cat#: A-11008 |  |
| Antibody | Rabbit Polyclonal anti-CAMSAP1L1 | Novus Biologicals | Cat#: NBP1-21402 | Antibody against CAMSAP2 |

### Mouse utilization

Ins-Apl mice with Histone 2B-mApple knocked into the Ins2 locus (Stancill et al., 2019) were typically used. Ins-Apl homozygous mice crossed with GFP-ELKS knock-in mice (Noordstra et al. 2022), were used for experiments with ELKS. Ins-Apl homozygous/GFP-ELKS mice crossed with homozygous Halo-Lifeact transgenic mice were used for experiments with Lifeact. The Halo-Lifeact transgenic mouse line was generated in the Vanderbilt Genome Editing Resource (VGER) and was generously gifted by Mathew Tyska. Males and females between 2 and 6 months were used. Mouse utilization was supervised by the Vanderbilt Institutional Animal Care and Use Committee (IACUC). Wild-type (WT) CD-1 (ICR) mice used in Western blotting were from Charles River Laboratories (Wilmington, MA).

### Islet Picking and Attachment

Mouse pancreatic islets were hand-picked following in situ collagenase perfusion and digestion as described in Ho et al. (2023a). Islets were allowed to recover for at least 1 hr in RPMI 1640 Media (Thermo Fisher Scientific, Cat# 11875093) supplemented with 10% FBS, 100 U/ml penicillin, and 0.1 µg/ml streptomycin in 5% CO2 at 37°C.

All coverslips and dishes were plasma cleaned and coated in placenta-derived human ECM (Corning, Cat#: 354237) which is composed of a mixture of laminin, collagen IV, and heparan sulfate proteoglycan, which serves as a reconstitution of vasculature ECM to recapitulate the beta cell interaction with the basement-membrane type ECM produced by endothelial cells of islet capillaries, for 10 min at 37°C.

Islets treated with lentivirus received virus 24 hrs prior to attachment. For attachment, 2-3 islets per 10 mm glass-bottomed dish (MatTek, cat# P35G-1.5-10-C) were placed in the center of the glass in 50 µl RPMI 1640 and transferred to 5% CO2 at 37°C. The following day, 50 µl of RPMI 1640 was added. Experiments were performed after 4-5 days of attachment. It has been previously shown that islets attached to vascular ECM preserve normal ability to secrete in response to glucose for up to 14 days (Patterson et al., 2000; Zhu et al., 2015).

### Cell Lines and Maintenance

MIN6 cells (Miyazaki et al., 1990) were maintained in DMEM with 10% fetal bovine serum (FBS) and 1% penicillin-streptomycin antibiotic at 37°C with 5% CO2 and were periodically tested for mycoplasma.

### Plasmids and Lentivirus

pcDNA3.1(+)/Puro/TagRFP-LL5b was a gift from Tomasz Prószyński & Joshua Sanes (Addgene plasmid # 112828 ; http://n2t.net/addgene:112828 ; RRID:Addgene_112828) and was cloned into a lentiviral backbone to produce pLV[Exp]-Puro-CMV>{TagRFP-LL5b} (RFP-LL5β) by VectorBuilder (Vector ID VB230831-1644ppw). TUBB5-Halo was a gift from Yasushi Okada (Addgene plasmid #64691; http://n2t.net/addgene:64691; RRID:Addgene_64691) and was cloned into a lentiviral backbone to produce pLV[Exp]-CMV>{TUBB5-Halo} (Halo-Tubulin) by VectorBuilder (Vector ID VB230330-1644hzc). pHR-scFv-GCN4-HaloTag-GB1-NLS-dWPRE (Halo-NLS) was a gift from James Zhe Liu (Addgene plasmid #106303; http://n2t.net/addgene:106303; RRID:Addgene_106303). pHRdSV40-K560-24xGCN4_v4 (SunTag-KIF5B) was a gift from Ron Vale (Addgene plasmid # 72229 ; http://n2t.net/addgene:72229 ; RRID:Addgene_72229). pEGFP-C1-CAMSAP2 (GFP-CAMSAP2) plasmid was a gift from Dr. Stephen Norris (Calico Life Sciences) and was cloned into a lentiviral vector in Ho et al. (2023). mCherry-UtrCH in the lentiviral backbone was generously gifted by Matthew Tyska.

### Lentivirus Production and Transduction

Lentivirus was produced by mixing the transfer plasmid with packaging plasmids pMDL, pRSV, and pVSV-G in a 6:4:1:1 ratio. The mixture was incubated in Optimem (Gibco) with TransIT-Lenti reagent (Mirus Bio) and subsequently added to 70% confluent HEK293 cells followed by 48h incubation at 37°C and 5% CO2. Supernatant containing lentivirus was precleared by pelleting cell debris through centrifugation and concentrated using Amicon Ultra-15 centrifugal filters (MilliporeSigma). Virus was stored at -80°C and added to islet media directly upon islet attachment.

### Western Blotting

Isolated mouse islets were lysed in Tris-lysis buffer (10mM Tris-HCl pH 75, 100mM NaCl, 1% Triton X-100, 10% glycerol, and complete protease inhibitor cocktail [Roche, Basel, Switzerland]) on ice and briefly sonicated. For SDS-PAGE, 30µL of lysate was loaded per lane and resolved in a 7% acrylamide gel. Gels were cut and probed using Rabbit anti-ERC1 for ELKS (1:1000, Atlas Antibodies), Mouse anti-GAPDH (1:5000, Santa Cruz Biotechnology), IRDye 680 Goat anti-Rabbit IgG (1:5000, LI-COR BioScience), and IRDye 800 Goat anti-Mouse IgG (1:7000, LI-COR BioScience). Blots were imaged using the Odyssey CLx imager (LI-COR).

### Microscopy

Fixed samples were imaged on a laser scanning confocal microscope Nikon A1r based on a TiE Motorized Inverted Microscope using a 100x lens, NA 1.49, run by NIS Elements C software. Cells and islets were imaged in 0.2 µm slices through the whole cell for MIN6 beta cells and through 10µm of the bottom of the islet. Select tubulin and Lifeact images were acquired using a Nikon Ax using the NSPARC detector system with a 100x lens, NA 1.49, with NIS Elements C.

FluoZin-3 assays and live intact islets were imaged on a Nikon TE2000E microscope equipped with a Nikon TIRF2 System for TE2000 using a TIRF 100x 1.49 NA oil immersion lens and an Andor iXon EMCCD camera run by NIS Elements C software.

Laser excitation wavelengths of 480, 560, and 640nm were used. Frame rate for Figs. 1, 3, 4, 5, S3, and S4 was 16 frames/second. Frame rate for Figs. 2 and S3 was 8.1 frames/second. Frame rate for Figure 6 was 1.1 frames/second.

Supplemental figure 1 was imaged on Zeiss LSM 980 microscope with Airyscan 2 detector, using 40x lens, NA 1.3, and processed by Zeiss Zen software.

### FluoZin-3 Secretion Assay

The FluoZin-3 assay (Gee et al., 2002; Zhu et al., 2015; Figure S2, Video 1.1) was performed 4-5 days after picking to allow for robust attachment of the islets. On the day of the assay, islets were incubated at 37°C in low glucose (2.8 mM glucose) KRB (110 mM NaCl, 5 mM KCl, 1 mM MgSO_4_, 1 mM KH_2_PO_4_, 1 mM HEPES, 2.5 mM, CaCl_2_, and 1 mg/ml BSA) for 2-4 hr with a change of buffer after 1 hr. For experiments using HaloTag 646 ligand (Promega, cat# GA1121), ligand was added to a final concentration of 800nM for at least 2 hrs. Immediately before imaging, the buffer was replaced with 100 µl fresh buffer to reduce background.

Dishes were placed on the TIRF microscope and allowed to equilibrate for at least 10 min. Islets were identified by eye. A 10 µm stack of 0.2 µm slices was recorded of the islet before addition of the FluoZin-3 dye using both transmitted light and the 568 nm laser to identify beta cells with red nuclei. Stacks were started below the islet to ensure the bottom of the cells were imaged. 38.8 µl of KRB buffer with the cell-impermeant FluoZin-3 dye (Thermo Fisher Scientific, Cat#: F24194) to final concentration of 20 µM dye was added. For high glucose treatment, glucose to a final concentration of 16.7 mM was added together with the dye.

Focus and TIRF angle were refined just prior to dye addition and the recording (30 ms exposure, no delay) was started within 30 sec after glucose stimulation to register active GSIS. The maximal recording time was restricted to 6-8 min to capture first phase secretion only and avoid photo-toxicity.

### VAEM Microscopy

For variable angle epifluorescence microscopy, the TIRF critical angle was first determined, followed by an angle which allowed for structures visibly not in the TIRF plane. For dual TIRF/VAEM imaging, triggered acquisition was used, in which the angle switched between TIRF and VAEM without delay. See Figure S5 for schematic.

### Analysis of FluoZin-3 Movies for Secretion Event Localization

FluoZin-3 assay analysis was performed as described in Trogden et al. (2021). Briefly, in ImageJ, the first frame of the movie was removed from one copy, and the last frame of the movie was removed from another copy. The movie copies were divided by each other using the Image Calculator tool in ImageJ with the 32-bit (float) result box checked. This resulted in a movie with all signal except for FluoZin-3 flashes removed. This processed movie was converted to an .h5 file for input into two custom Ilastik machine learning programs for 1) pixel classification and 2) object classification. Center point coordinates obtained from object classification were used for analysis.

Cells were identified using transmitted light stacks to determine cell borders and beta cell identity. All cells with a red nucleus (Ins-Apl signal) were outlined by hand in ImageJ. If the nuclei could not be seen (above the image stack range, signal diminished because of light dispersal or out of the frame) or was Ins-Apl negative, the cell was outlined but assumed to be a non-beta cell. Each Ins-Apl positive cell outline was saved as an individual ROI in ImageJ and coordinates were exported.

Secretion event and cell outline coordinates were imported into a custom MATLAB script as in Trogden et al. (2021). The script uses density-based scanning on a point-by-point basis to identify secretion events that have a minimum of three neighbors within a 1.5µm radius. Secretion events meeting these criteria are identified as a cluster. Analysis output included events per cell, clustered events, coordinates, cluster ID, and counts per cluster.

#### Analysis of Secretion at ELKS and LL5β patches

Knock-in GFP-ELKS islets were used with the FluoZin-3 assay. ELKS patches were manually thresholded on a 0.7 Gaussian filter in ImageJ for blurring to generate a more inclusive patch threshold. Secretion events were identified as described above. The first frame of the movie was used to create ELKS thresholds/masks, due to the fact that FluoZin-3 will eventually saturate the ELKS signal in green and prevent precise live ELKS tracking. Prior to this analysis, we confirmed that we observed little net ELKS movement over the course of 6-8 minutes of imaging. This technique was applied to other green fluorophore probes.

The total ELKS area was also measured in µm^2^, and secretion events were treated as 0.23µm^2^ occurrences, a size chosen for diffraction limit, approximate pore size, secretion machinery size, etc. Total secretion event area was calculated as a percentage of total ELKS area.

The same analysis was performed with RFP-LL5β-transduced islets, except that cells not expressing RFP-LL5β were excluded from analysis due to inability to reconcile endogenous LL5β and labeled LL5β. Ins2Apl-expressing beta cell nuclei were manually excluded from RFP-LL5β threshold creation.

#### ELKS intensity Analysis for Distribution Histograms

Intensity analysis was performed for ELKS using a custom macro for exporting mean gray value of each pixel from a non-rectangular ROI in ImageJ. The mean gray value of ELKS at the secretion event pixel, of every pixel within ELKS patches, and of every pixel within the whole cell was obtained for each cell. A rectangular region outside of the ELKS signal was taken to be background and mean gray value was measured. Background was subtracted from all values, and all values were divided by the highest pixel value in the cell to obtain a 0-1 scale. This normalization was performed for each cell in each islet to compensate for the slight illumination differences in TIRF microscopy, to best normalize for each cell.

#### Analysis of MT Minus Ends

Islets were transduced with GFP-CAMSAP2 lentivirus at a 1:50 dilution for 24h before plating. Images were processed as described above, and MT minus ends indicated by CAMSAP2 were manually identified in ImageJ. If the CAMSAP2 was punctate, only one end was chosen. If the CAMSAP2 decorated a stretch of a MT, both ends were chosen, as we cannot say with certainty which end is the actual minus end. An ImageJ macro for determining shortest distance between one point and others (https://microscopynotes.com/imagej/shortest_distance_to_line/distance_macros_v102.txt) was used to assess shortest distance between each secretion event and each CAMSAP2 puncta/stretch.

#### Analysis of MT Plus Ends

For assessing MT directionality and plus ends, islets were transduced with SunTag-KIF5B with Halo-NLS at a 1:20 and 1:50 dilution, respectively. Islets were treated with HaloTag ligand prior to experiments (see above). Images were processed as described above, individual cells were masked out for ease of analysis, and KIF5B puncta were tracked using Imaris (Oxford instruments, v10.1) tracking software. The shortest distance macro was applied to assess shortest distance between secretion events and KIF5B track enpdoints. A custom MATLAB script previously developed in and used by our lab (Trogden et al., 2021) was used to assess clustering of KIF5B track endpoints and secretion events, respectively. Clusters were visualized using the makePolygon function in ImageJ. The shortest distance macro in ImageJ was then applied between KIF5B track endpoint clusters and secretion clusters. Track displacement data were also exported from Imaris, and compared between low and high glucose-treated islets.

#### Analysis of MT Configuration at Secretion Event Location

Movies acquired by semi-simultaneous TIRF/VAEM imaging were blinded and analyzed manually by two individuals. GFP-ELKS islets were transduced with Halo-Tubulin lentivirus at a 1:50 dilution for 24h before plating. Islets were treated with HaloTag ligand prior to experiments (see above). Images were processed as described above and ELKS and MTs were analyzed in ImageJ for the following: 1) orientation of MTs relative to secretion events; 2) Orientation of MTs prior to secretion events when 1) was N/A; and 3) Shortest distance between MTs and secretion events when 1) was N/A. Shortest distances were measured as described above.

### Immunofluorescence

For CAMSAP2 staining, MIN6 cells transduced with GFP-CAMSAP2 or not (endogenous only) were plated on fibronectin-coated coverslips and allowed to attach for 24 hours. Cells were then fixed in ice cold anhydrous methanol for 5 minutes followed by three washes in PBS. Cells were permeabilized in 0.1% Triton-X100 in PBS for 15 minutes, followed by three washes in PBS. Cells were blocked using BSA/DHS for 1 hour, washed three times in PBS, then incubated in primary antibody for 4h at room temperature or 24h at 4°C. Following three washes in PBS, cells were incubated in secondary antibody for 4h at room temperature or 24h at 4°C. After three final PBS washes, coverslips were mounted on slides.

For LL5β staining in intact mouse islets, islets attached to vascular ECM as described above were fixed in 4% paraformaldehyde with 0.1% saponin and 0.3% Triton X-100 for 1 hour, followed by three washes in PBS containing 0.1% saponin/Triton X-100. Cells were blocked using BSA/DHS with 0.1% saponin/Triton X-100 for 2 hours, washed three times in PBS, then incubated in primary antibody with 0.1% saponin/Triton X-100 overnight at 4°C. Following three washes in PBS, cells were incubated in secondary antibody with 0.1% saponin/Triton X-100 for 4h at room temperature. After three final PBS washes, coverslips were mounted on slides.

Antibodies used included Rabbit anti-CAMSAP1L1 (1:250, Novus Biologicals), Goat anti-Rabbit 488 (1:500 for CAMSAP2, 1:200 for LL5β), andRabbit anti-PHLDB2 (1:200) for staining, and anti-GFP-FITC for labeling GFP-CAMSAP2-transduced cells. Coverslips were mounted in VectaShield mounting medium (Vector Labs, Cat#: H-1000).

### Statistics

For all data sets unless otherwise noted, Student’s t test was used with P<0.05 for significance. Graphical representations and statistical analysis were performed using GraphPad Prism.

### Experimental Design

All experiments were replicated at least three times, from at least three different mice for biological replicates, on at least three different days. At least 9 islets per experiment were imaged for three technical replicates per biological replicate. For all GFP-ELKS islets, all cells were analyzed. For all RFP-LL5β islets, cells expressing RFP-LL5β were analyzed and those not expressing were excluded from analysis. For MT end analysis, cells expressing SunTag-KIF5B, GFP-CAMSAP2, or Halo-Tubulin were analyzed and those not expressing were excluded from analysis unless otherwise stated for internal control purposes.

For all islets, if at least one secretion event was recorded, it was analyzed. For islets not displaying secretion, these were either considered “zero” for secretion purposes, or excluded and determined to be internal controls as they could not be factored into analysis if they were low glucose-treated. Analysis of MT configuration was performed double blinded.

### Image processing

The majority of imaging data are single-slice TIRF or VAEM images. Confocal data in Figure 5A shown as maximum intensity projection, in Figure 5B shown as single slice. Airyscan images are presented as single XY optical slices and YZ projection produced in ImageJ. All images have been pseudo-colored for consistency of presentation. Histogram stretching was applied to most whole images for better visualization. In Figs. 6, S1, S2, S5, and S6, slight adjustments were made to gamma settings to make small structures visible. For Videos 3.1-3.3, FluoZin-3 flashes were blurred using a 2.0 Gaussian filter in ImageJ for clarity. For videos with FluoZin-3 flashes, grouped Z projections were used to clarify flashes.

## Acknowledgements

This work was supported by National Institutes of Health (NIH) grants F31 DK13344-01 and T32 DK007563-33 (to MAF; O’Brien, PI), NIDDK R01 DK106228 MPI (to IK and GG), NIGMS MIRA R35GM127098 (to IK), R01-DK65949 (to GG), and R01-DK125696 (to GG). PN was supported by by an NIH training grant T32 DK101003 (Young, PI). We thank Hamida Ahmed for technical help. We thank Zeiss for Airyscan microscopy demonstration. We thank the Diabetes Resesarch Training Center (DRTC) and members for advice and support. We also thank all members of the Kaverina laboratory and the Vanderbilt Mechanobiology & More Club for advice and support. Schematics were made using Biorender.

## Abbreviations

MT: Microtubules
IG: insulin granules
TIRFM: total internal reflection fluorescence microscopy
CMSC: cortical microtubule stabilizing complex
ECM: extracellular matrix
VAEM: variable angle epifluorescence microscopy

