## Supplementary Material for "Directed insulin secretion from beta cells occurs at cortical sites devoid of microtubules at the edges of ELKS/LL5β patches"

**Supplemental Figure Legends**

**Figure S1.** Expression levels in GFP-ELKS knock-in mice and RFP-LL5β-transduced islet cells. A) Western blot of endogenous ELKS (WT) compared to GFP-ELKS knock-in protein levels with GAPDH loading control. B) Quantification of A) normalized to total protein content. N=3 mice per group. C) Secretion events per islet for WT and GFP-ELKS islets. N=13 islets per group, n=59-267 secretion events for WT, n=37-231 secretion events for GFP-ELKS. D-F) Comparison of endogenous and ectopic RFP-LL5β intensity levels. D) Endogenous LL5β. Outlines of cells not expressing RFP-LL5β. E) Ectopic RFP-LL5β. Outlines of cells expressing RFP-LL5β. F) Composite of D-E). G) Relative intensity of endogenous LL5β in cells expressing RFP-LL5β or not. N=8 islets, n=34 cells per group. Student's t test, $P<0.05$ for all.

**Figure S2.** Experimental system using intact mouse islets shows that GFP-ELKS primarily localizes to the bottom of the islet, at the point of attachment to vascular ECM. A) Schematic of the islet attachment to a MatTek dish coated with vascular ECM. B) XZ (side) view of a GFP-ELKS (magenta) and mApl-Ins2 (green nuclei) in a mouse islet, attached to vascular ECM. Cyan arrowhead, lower optical plane at the ECM attachment level, corresponding to C). Yellow arrowhead, higher optical plane deeper in the islet, corresponding to D). White lines, sectioning for C). Scale bar = 10µm. C) Lower optical plane, showing a high localization of GFP-ELKS at the vascular ECM/attachment site. Scale bar = 10µm. D) Higher optical plane, showing GFP-ELKS accumulation in smaller patches, presumably around vascular structures maintained within the islet (arrows). Note that the GFP-ELKS intensity at vascular structures within the islet is similar to the intensity of ELKS patches at the vascular ECM-coated coverslip. E-G') Method of identifying beta cell outlines and nuclei. E) Brightfield image of intact islet with E') cell outlines. F) Ins-Apl labeled beta cell nuclei with F') cell outlines. G) Composite of E-F with G') cell outlines.

**Figure S3.** Focal adhesions approximated by Halo-Lifeact or mCh-UtrCH localize adjacent to ELKS puncta. A) Islet beta cell expressing knock-in GFP-ELKS and transgenic Halo-Lifeact,maxIP of the bottom 1µm of the cell using NSPARC super-resolution microscopy. B) Inset shows an ELKS puncta adjacent but not overlapping the end of an actin filament (white arrow). C-F) Islet beta cells expressing knock-in GFP-ELKS and ectopic mCh-UtrCH. C) NSPARC of bottom 1μm of the cell with D) inset showing ELKS adjacent to actin filament end (white arrow). E) TIRF plane of cell with F) inset.

**Figure S4.** Experimental system using the FluoZin-3 assay allows for precise spatial and temporal localization of secretion events. A) Schematic detailing the experimental setup. B) Schematic detailing TIRF method, with fluorophores in the TIRF plane. C) Example output of the MATLAB script, showing xy coordinates in a 512x512 pixel image. D-K) Montage of a secretion event. At D) 0ms there is no secretion event visible, at E) 100ms the flash starts to appear, at F) 200ms the flash is easily visible, and at G) 300ms, the flash has disappeared again. H-K), same as D-G), with secretion event xy coordinate identified in I-J) (green cross). Corresponds to Video 1.1. L-M) High-confidence, smaller radius of Ilastik-identified (white outline) secretion event (green cross). N-O) Lower-confidence, larger radius of Ilastik-identified secretion event. P) Time projection of a FluoZin-3 movie displaying time localization of secretion events.

**Figure S5.** Line scans of FluoZin-3-detected secretion events as compared to ELKS intensity. A-C) Line scan of an event at high-intensity ELKS, corresponding to Fig. 3. A) One frame before secretion event, B) secretion event and two lines for multiple comparisons, C) one frame after event. D) Line scan of Line 1, E) Line scan of Line 2. F-H) Line scan of an event at low-intensity ELKS. I-J) Line scan plots for low-intensity. K-M) Line scan of an event occurring outside of ELKS. N-O) Line scan plots for zero-intensity.

**Figure S6.** ELKS and LL5β time projections reveal minimal dynamics over the acquisition period. A) Intact islet expressing GFP-ELKS with no FluoZin-3 treatment to allow for clear visualization. Time projection with B) inset. C) Intact islet expressing RFP-LL5β with D) inset.

**Figure S7.** GFP-ELKS and RFP-LL5β co-localize in mouse islets in cortical patches that also display secretory heterogeneity. A) Islet cell outlines (white) with insulin secretion events (yellow crosses), and non-beta cells (red). Cells not expressing RFP-LL5β, gray. Single-plane TIRF images showing an intact mouse islet expressing B) knock-in GFP-ELKS (grayscale) or C) ectopic RFP-LL5β (grayscale), forming patches at the interface with vascular ECM. Scale bar = 10μm. D) Threshold of GFP-ELKS (magenta) and RFP-LL5β (green) patches. E) Inset of A). Insets of F) GFP-ELKS (magenta) and G) RFP-LL5β (green) patches. Scale bar = 1μm. H) Composite of GFP-ELKS (magenta) and RFP-LL5β (green) with insulin secretion events (yellow crosses). I) Composite of GFP-ELKS (magenta) and G) RFP-LL5β (green) with corresponding color outlines. Thresholds of J) GFP-ELKS (magenta) and K) RFP-LL5β (green). L) Composite of thresholds, with insulin secretion events (gray crosses). Gray mask excludes cells and events that are excluded from analysis, according to A).

**Figure S8.** An example of MT minus end analysis in a cell with CAMSAP2 stretches. A) GFP-CAMSAP2 expressed ectopically in intact mouse islets. Higher CAMSAP2 expression levels results in decoration of the MT in a stretch pattern. In this case, both ends were considered MT minus ends (see Materials & Methods). Scale bar = 1μm. B) Previous panel, with secretion events (yellow crosses). C) Previous panel, with CAMSAP2 stretch ends (cyan). D) Previous panel, with example of shortest distance measurements (magenta).

**Figure S9.** VAEM and TIRF combination allows for full MT visualization. A) Halo-Tubulin in the TIRF plane. Scale bar = 1μm. B) Halo-Tubulin in the VAEM plane. C) Composite image showing the anchoring points of MTs in TIRF plane in both end-on (arrowhead) and lattice-bound (arrow) conformations. D) Schematic of C), showing an end-on MT with its end in the TIRF plane (red glow, arrowhead) and a lattice-bound MT with its anchoring point in the TIRF plane (red glow, arrow).

**Figure S10.** Three distinct MT conformations exist in beta cells. A) End-on configuration, showing Halo-Tubulin in TIRF (red) and VAEM (cyan). Arrows, anchoring points of MTs in TIRF. Scale bar = 1μm. B) Lattice-bound configuration. C) No association with GFP-ELKS. D) End-on configuration, showing Halo-Tubulin in VAEM (cyan) anchored at GFP-ELKS (magenta). Arrows, anchoring points at GFP-ELKS. E) Lattice-bound configuration. F) No association with GFP-

ELKS. Arrow, GFP-ELKS patch without MTs associated. G) End-on schematic. H) Lattice-bound schematic. I) No association schematic. J-L”) Second example of MT withdrawal prior to insulin secretion event, with Fig. 6K-M”).

**Video Legends**

**Video 1.1**- with Figure S1

Insulin secretion in intact mouse islet using FluoZin-3, showing an individual secretion event as a flash. The flash is visible in 2 subsequent frames in this example. FluoZin-3, grayscale. See Materials & Methods for background subtraction protocol. Time after high glucose stimulation in seconds and milliseconds is shown.

**Video 1.2**- with Figure 1

Insulin secretion from a GFP-ELKS knock-in mouse islet, showing secretory heterogeneity. Magenta, GFP-ELKS, a still frame from the beginning of the recording. Green, secretion events over time via FluoZin-3, after background subtraction. Arrowhead = an actively secreting ELKS patch. Arrow = a dormant ELKS patch, not secreting. Each frame is averaged over time (10 frames = 0.6 seconds). Time after high glucose stimulation in minutes and seconds is shown.

**Video 2**- with Figure 2

Insulin secretion from RFP-LL5β expressing cells in a mouse islet (inset same as main figure), showing a secreting LL5β patch. Green, RFP-LL5β. Magenta, secretion events via FluoZin-3 after background subtraction. Each frame is averaged over time (10 frames = 1.1 seconds). Time after high glucose stimulation in minutes and seconds is shown.

**Video 3.1**- with Figure 3C

Example of an insulin secretion event (FluoZin-3 after background subtraction, green) occurring at the margin of a high-intensity GFP-ELKS patch (magenta), corresponding to Fig. 3C. Left, GFP-ELKS still frame; middle, secretion event via FluoZin-3; right, overlay with secretion event labeled as yellow cross. Time after high glucose stimulation in minutes, seconds, and milliseconds is shown.

**Video 3.2**- with Figure 3D

Example of an insulin secretion event (FluoZin-3 after background subtraction, green) occurring at a low-intensity region of a GFP-ELKS patch (magenta), corresponding to Fig. 3D. Left, GFP-ELKS still frame; middle, secretion event via FluoZin-3; right, overlay with secretion event labeled as yellow cross. Time after high glucose stimulation in minutes, seconds, and milliseconds is shown.

**Video 3.3**- with Figure 3E

Example of an insulin secretion event (FluoZin-3 after background subtraction, green) occurring in a GFP-ELKS-negative area in the vicinity of a GFP-ELKS patch (magenta) corresponding to Fig. 3E. Left, GFP-ELKS still frame; middle, secretion event via FluoZin-3; right, overlay with secretion event labeled as yellow cross. Time after high glucose stimulation in minutes, seconds, and milliseconds is shown.

**Video 4**- with Figure 4

Examples of high-displacement and low-displacement SunTag-KIF5B (grayscale) tracks. Upper track, high displacement. Lower track, low displacement; time-coded. Track time is color-coded. Time after high glucose stimulation in minutes and seconds is shown.

**Video 5**- with Figure 5

Insulin secretion from GFP-CAMSAP2 expressing cell in a mouse islet showing secretion relative to CAMSAP2. Grayscale, GFP-CAMSAP2. Green, secretion events via FluoZin-3 after background subtraction. Each frame is averaged over time (10 frames = 0.6 seconds). Time after high glucose stimulation in minutes and seconds is shown.

**Video 6.1**- with Figure 6

Example of MT loss from a future site of secretion. The movie is paused at 01:35 min:sec and 01:53 to demonstrate MTs likely sliding away from the future site of secretion. Arrow indicates the shortening MT. At time of secretion (02:03 min:sec), MTs are absent from the secretion site (circled yellow). Red, Halo-tubulin visualized by TIRF microscopy. Cyan, Halo-Tubulin visualized by VAEM. Time after high glucose stimulation in minutes and seconds is shown.

**Video 6.2**- with Figure 6

Example of MT loss from a future site of secretion. The movie is paused at 03:35 min:sec and 04:28 min:sec to demonstrate MT disassembly or sliding away from the future site of secretion. Arrow indicates the shortening MT. At time of secretion (04:51 min:sec), MTs are absent from the secretion site (circled yellow). Red, Halo-Tubulin visualized by TIRF microscopy. Cyan, Halo-Tubulin visualized by VAEM. Time after high glucose stimulation in minutes and seconds is shown.

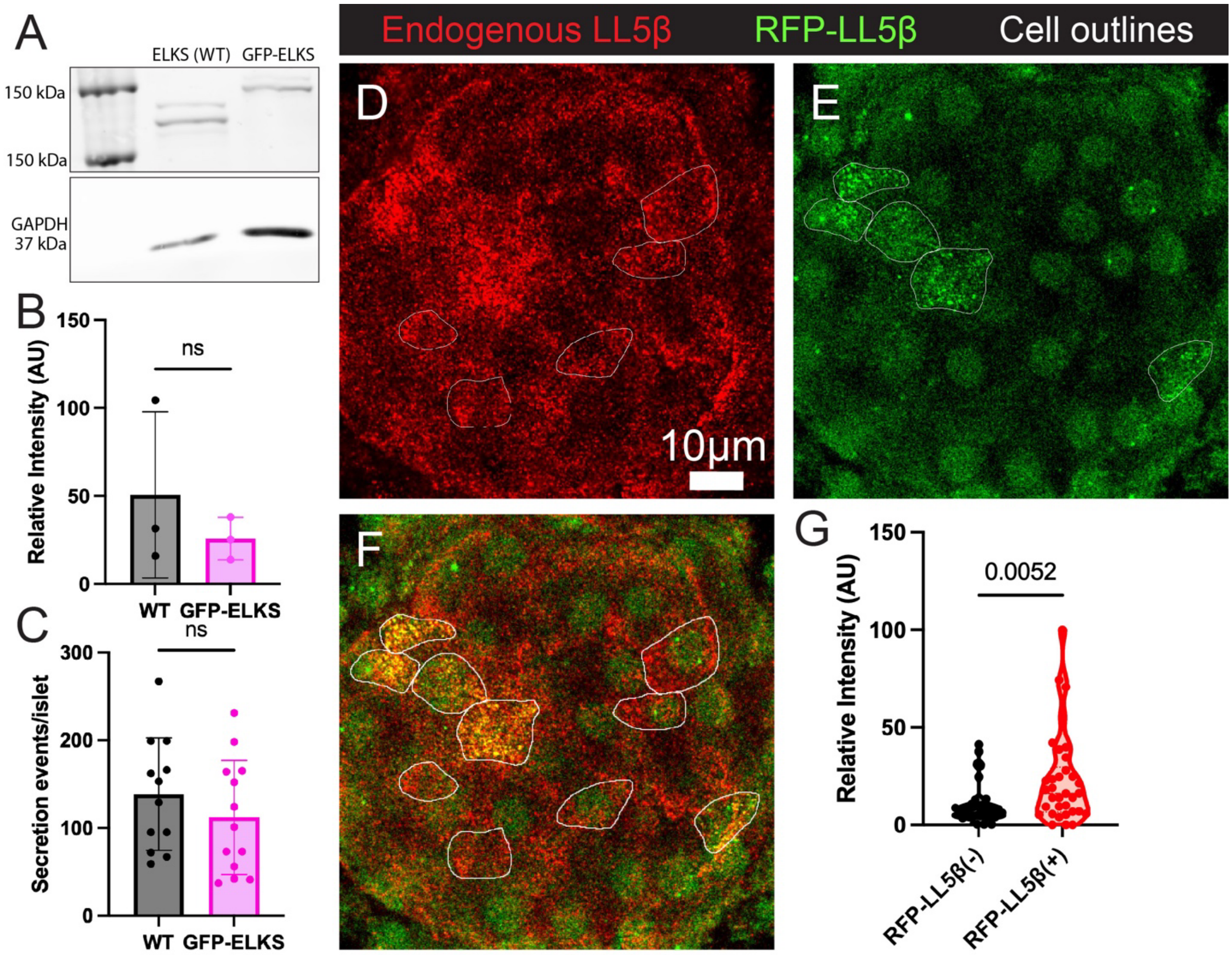
A
ELKS (WT)
GFP-ELKS
150 kDa
150 kDa
GAPDH
37 kDa
B
Relative Intensity (AU)
ns
WT
GFP-ELKS
C
Secretion events/islet
ns
WT
GFP-ELKS
Endogenous LL5β
RFP-LL5β
Cell outlines
D
E
10μm
F
G
Relative Intensity (AU)
0.0052
RFP-LL5β(-)
RFP-LL5β(+)

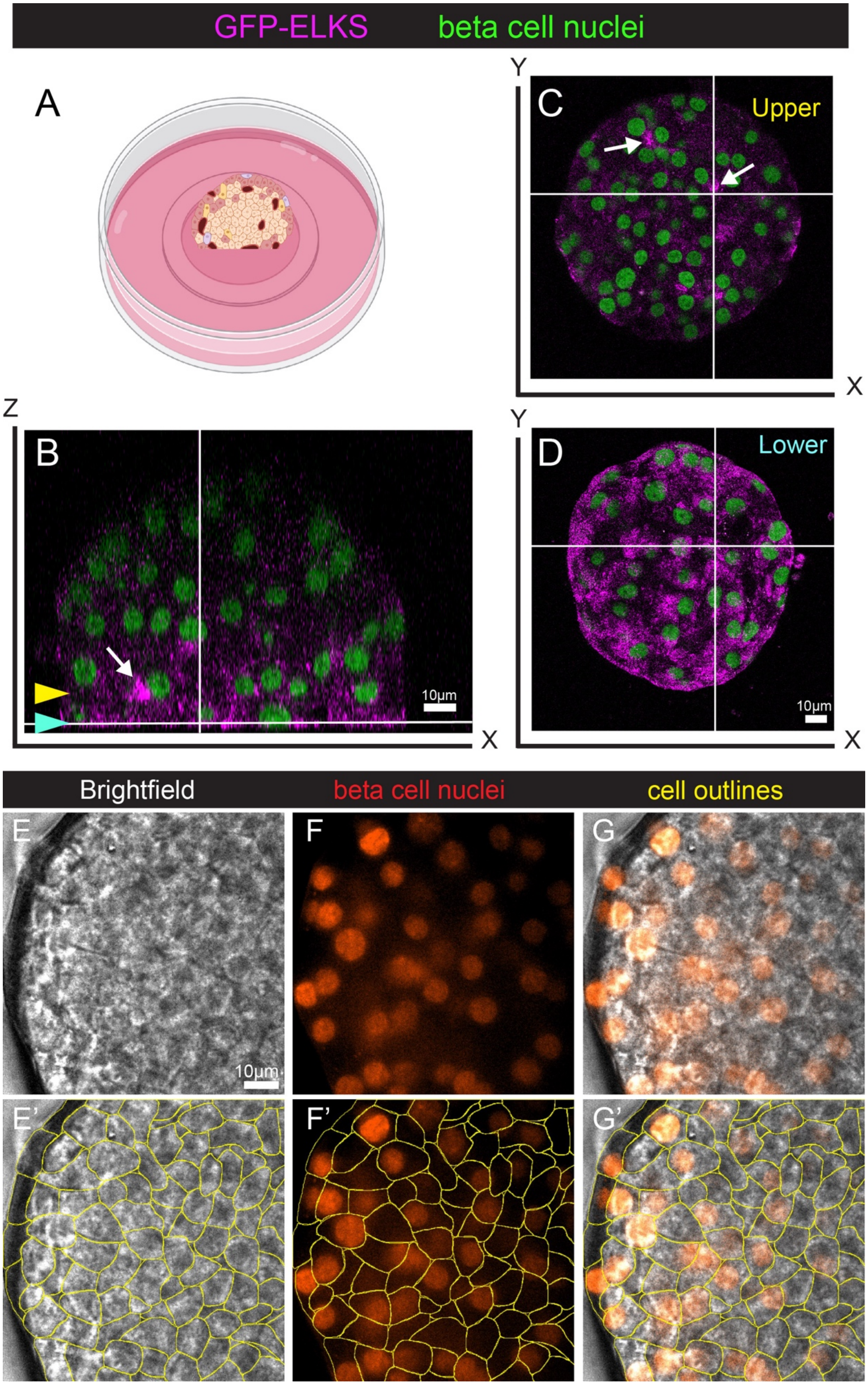

GFP-ELKS
beta cell nuclei
A
Y
C
Upper
X
Z
B
10μm
X
Y
D
Lower
10μm
X
Brightfield
beta cell nuclei
cell outlines
E
10μm
F
G
E'
F'
G'

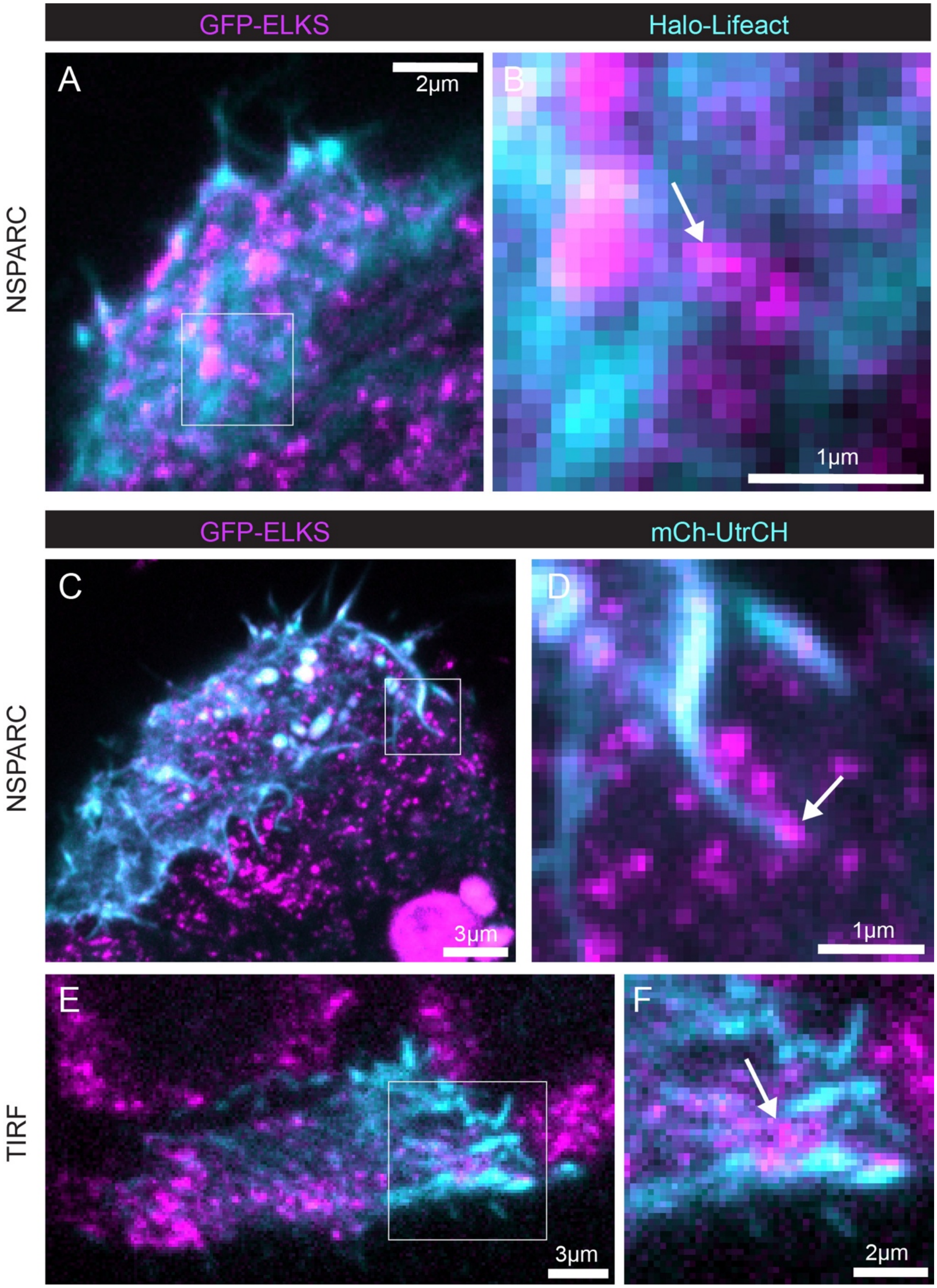

GFP-ELKS
Halo-Lifeact
NSPARC
A
2μm
B
1μm
GFP-ELKS
mCh-UtrCH
NSPARC
C
3μm
D
1μm
TIRF
E
3μm
F
2μm

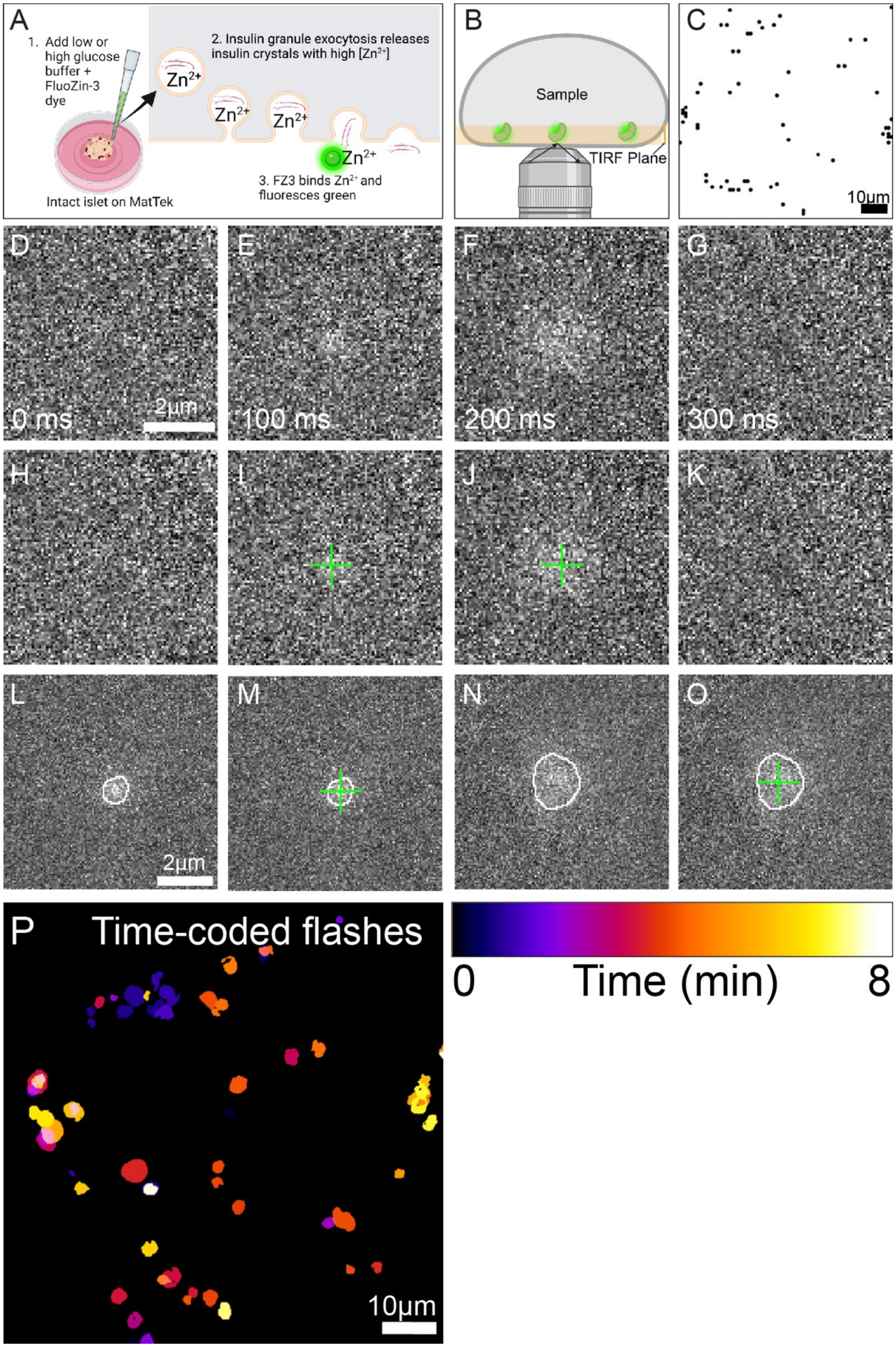
A
1. Add low or high glucose buffer + FluoZin-3 dye
Intact islet on MatTek
2. Insulin granule exocytosis releases insulin crystals with high [Zn2+]
3. FZ3 binds Zn2+ and fluoresces green
B
Sample
TIRF Plane
C
10μm
D
0 ms
2μm
E
100 ms
F
200 ms
G
300 ms
H
I
J
K
L
2μm
M
N
O
P
Time-coded flashes
10μm
0
Time (min)
8

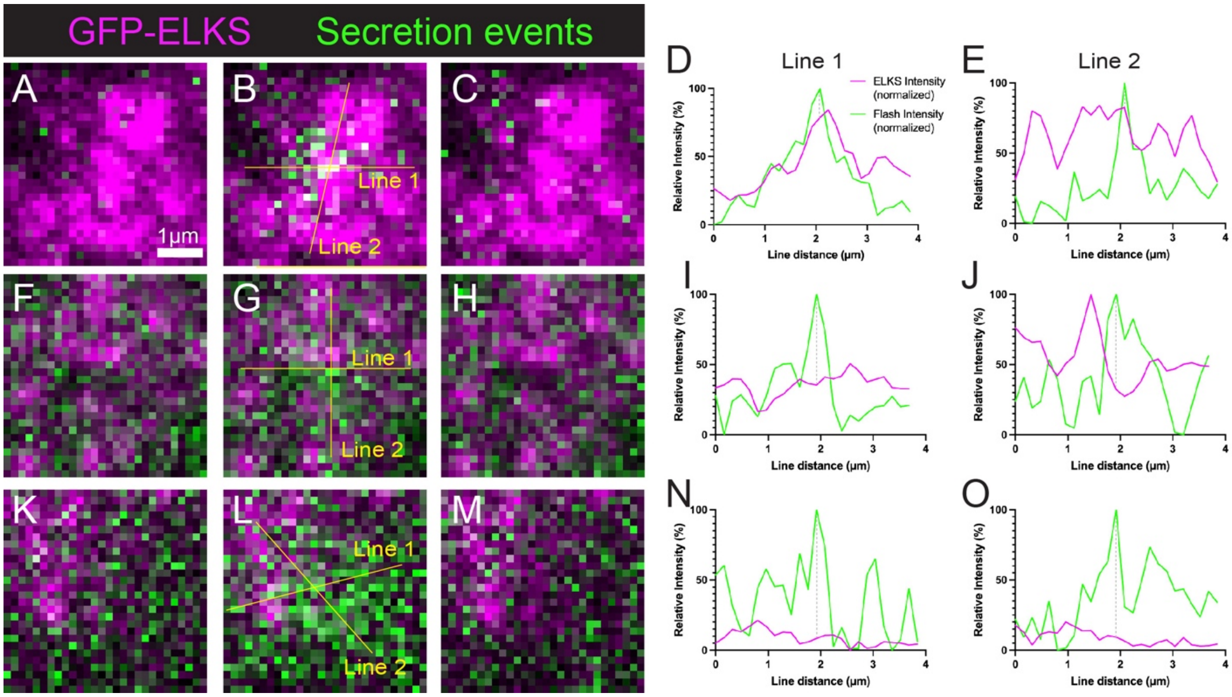

GFP-ELKS
Secretion events
A
B
C
1μm
Line 1
Line 2
D
E
Line 1
Line 2
ELKS Intensity (normalized)
Flash Intensity (normalized)
Relative Intensity (%)
Line distance (μm)
F
G
H
I
J
K
L
M
N
O

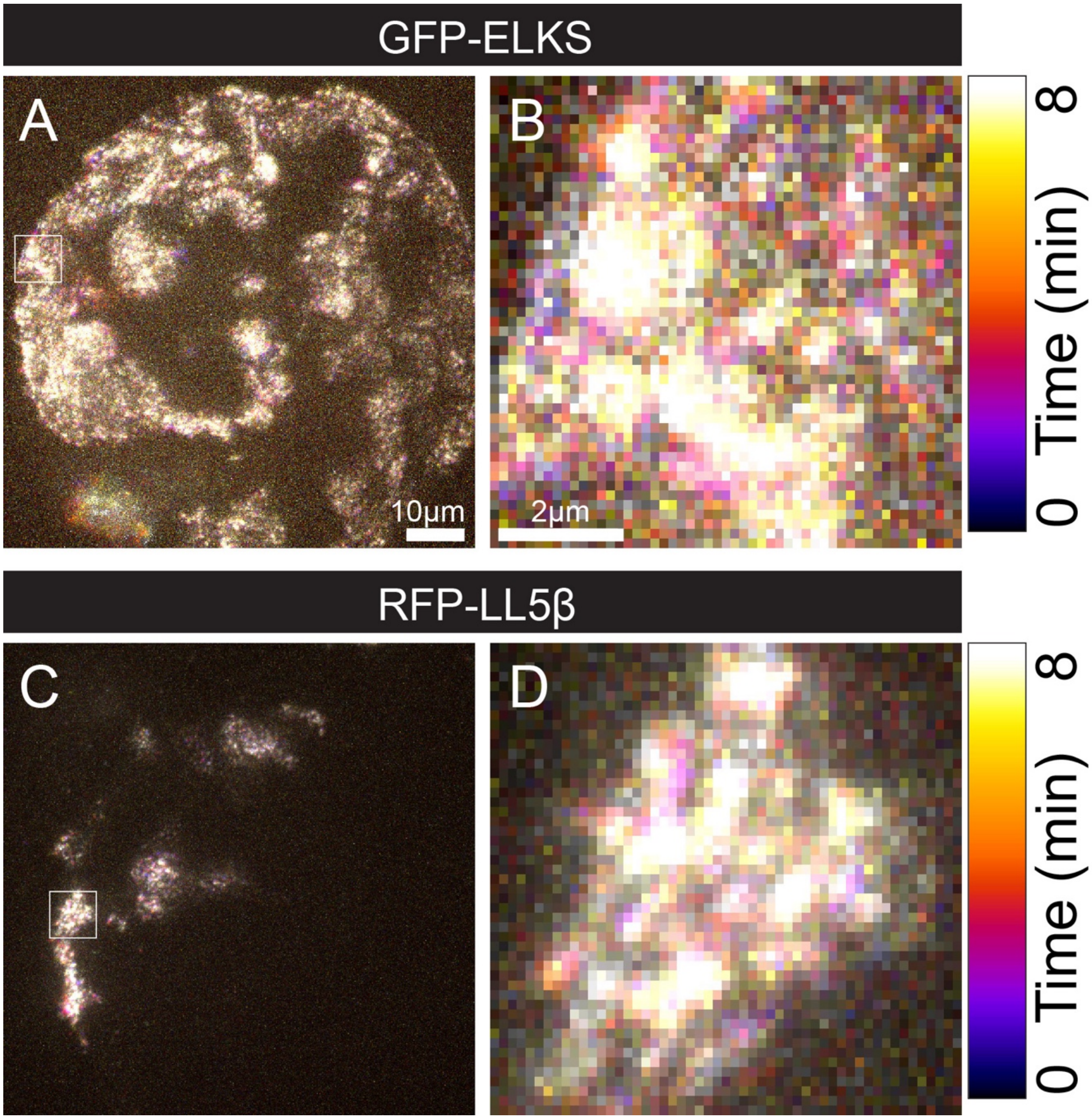

GFP-ELKS
A
B
10µm
2µm
8
Time (min)
0
RFP-LL5β
C
D
8
Time (min)
0

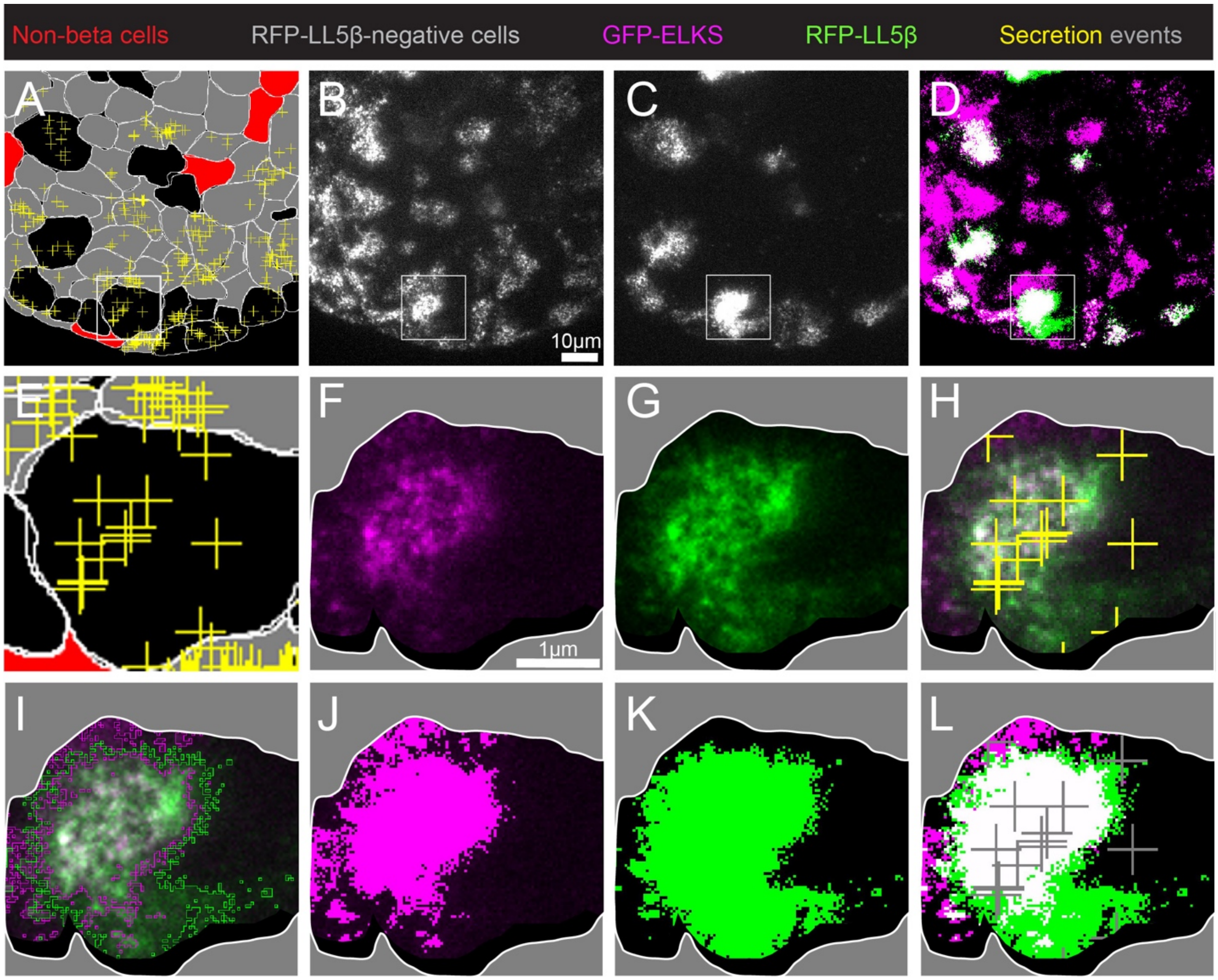

Non-beta cells
RFP-LL5β-negative cells
GFP-ELKS
RFP-LL5β
Secretion events
A
B
C
D
10μm
E
F
G
H
1μm
I
J
K
L

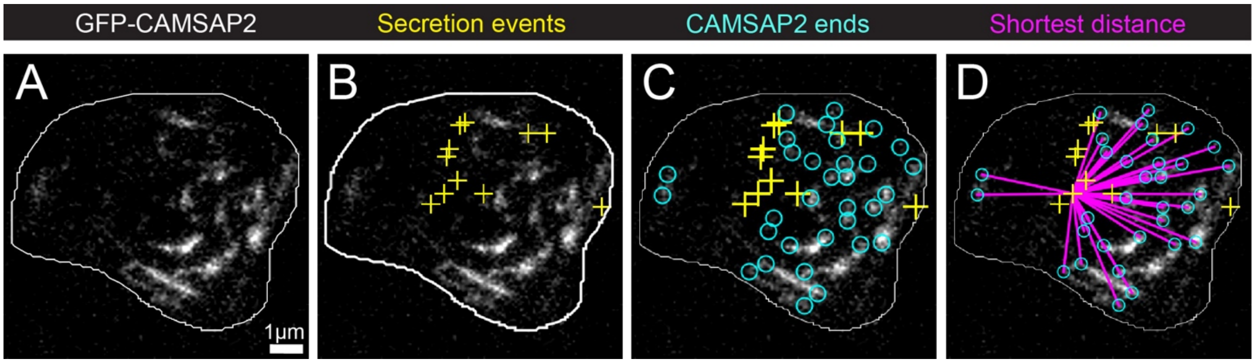
GFP-CAMSAP2
Secretion events
CAMSAP2 ends
Shortest distance
A
B
C
D
1µm

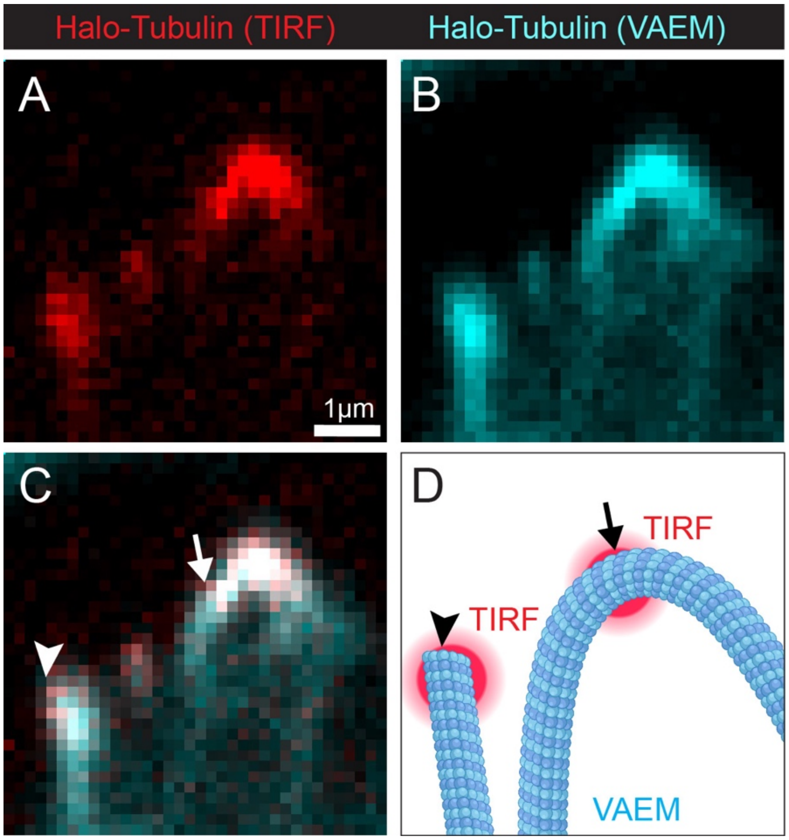

Halo-Tubulin (TIRF)
Halo-Tubulin (VAEM)
A
B
1μm
C
D
TIRF
TIRF
VAEM

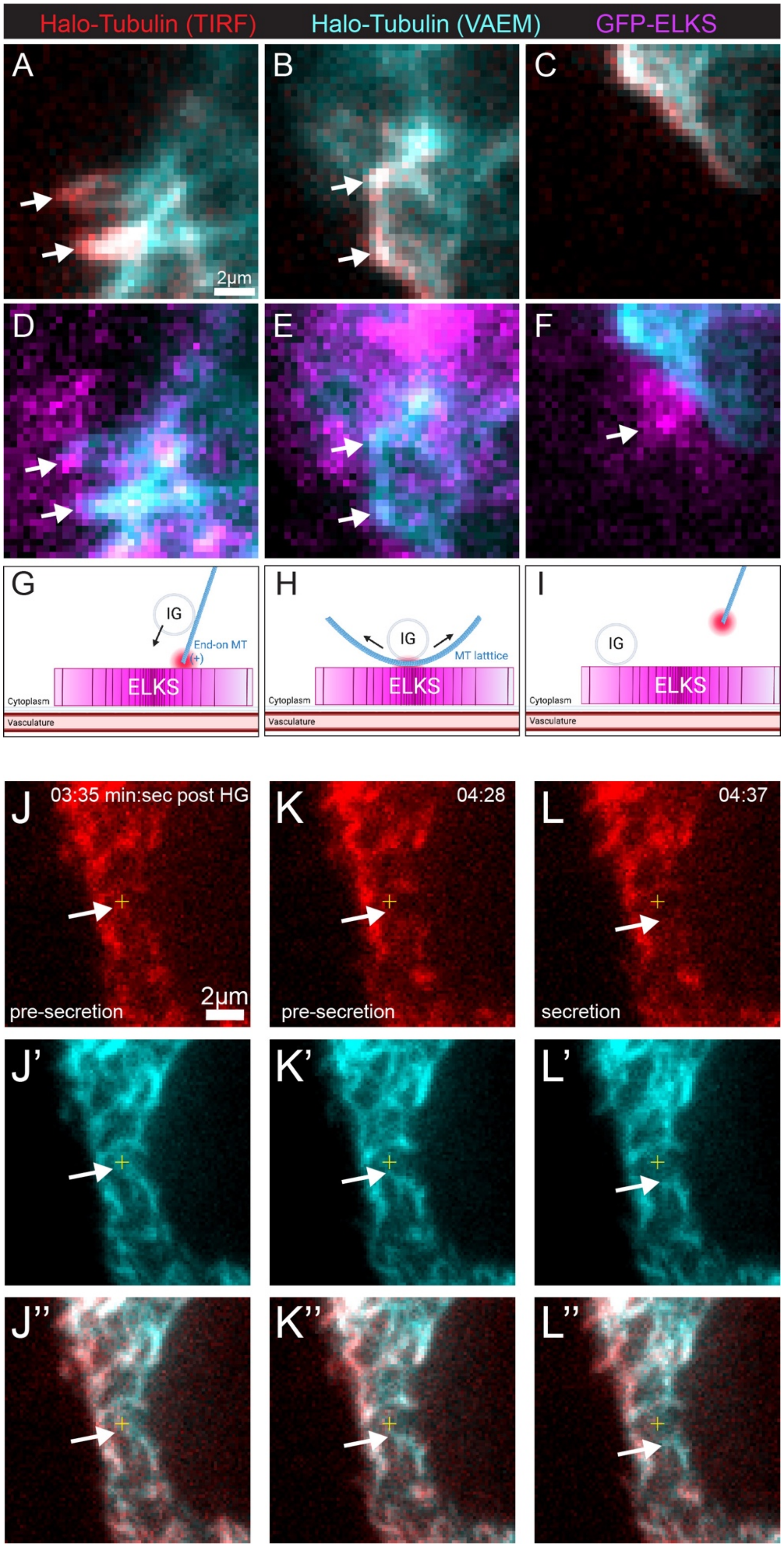

Halo-Tubulin (TIRF)
Halo-Tubulin (VAEM)
GFP-ELKS
A
B
C
2µm
D
E
F
G
IG
End-on MT
(+)
ELKS
Cytoplasm
Vasculature
H
IG
MT latttice
ELKS
Cytoplasm
Vasculature
I
IG
ELKS
Cytoplasm
Vasculature
J
03:35 min:sec post HG
pre-secretion
2µm
K
04:28
pre-secretion
L
04:37
secretion
J'
K'
L'
J''
K''
L''
